# Baldur: Bayesian hierarchical modeling for label-free proteomics exploiting gamma dependent mean-variance trends

**DOI:** 10.1101/2023.05.11.540411

**Authors:** Philip Berg, George Popescu

## Abstract

Due to its simplicity in sample preparation, label-free quantification has become de facto in proteomics research at the expense of precision. We propose a Bayesian hierarchical decision model to test for differences in means between conditions for proteins, peptides, and post-translation modifications. We introduce a novel Bayesian regression model to characterize local mean-variance trends in the data to describe measurement uncertainty and to estimate the decision model hyperparameters. Our model vastly improves over state-of-the-art methods (Limma-Trend and t-test) in several spike-in datasets by having competitive performance in detecting true positives while showing superiority by greatly reducing false positives.

## Background

Label-free quantification (LFQ) is a fast-growing methodology to infer abundances in mass spectrometry proteomics (1– 3). While a common issue in LFQ proteomics data is missing values (outside of the scope of this paper), it also tends to produce noisier data than labeling-based methods (4). Extensive research has focused on spectral quantification (2, 5–10) and identification (9–11). On the other hand, research towards understanding the final quantitative proteomics datasets and how to use data-set level information in statistical testing of differences in means is scarce. Mainly ANOVA or t-test methods are applied for this analysis (9, 11–14), but some work towards using mixed effect-, regression-models, and Stouffer’s method, have been developed for total proteomics analysis (15–18). Still, none of them can deal with generic datasets, e.g., peptidomics, phosphoproteomics, etc. Therefore, our focus here is on the statistical decision of the differences in mean abundances of peptides (or proteins, post-translation modifications, etc.) between different conditions (Wild-Type/mutant, control/treatment, time series, etc.). To this end, we present a Bayesian decision method (Baldur) that uses gamma regression to construct priors according to the mean-variance (M-V) of data and the uncertainty of individual measurements. In particular, we propose a new method for modeling the variance component using a gamma distribution. In addition, we develop an improved gamma regression model that describes the M-V dependency as a mixture of a common and a latent trend—allowing for localized estimates of the M-V trend. This is then used for inference of measurement uncertainty and hyperparameters for the variance prior. We then evaluate the performance of Baldur, t-test, and Limma-Trend on three total proteomics datasets and one post-translational modification (PTM) dataset. Importantly, we find that Baldur drastically improves the decision in noisier PTM data (19) over Limma-Trend and t-test. Likewise, we see significant improvements using Baldur over the other methods at small spike-in quantities in one of the total proteomics datasets (14). Further, we show that Baldur improves the performance on the remaining two total proteomics data (19, 20) over all other models by reducing the number of false positives. Lastly, we performed empirical power analysis to analyze the methods’ precision when increasing the number of replicates. We found that the Baldur methods always gain power with sample size while showing robust control of the false positive. On the other hand, Limma-Trend and t-test increase in power with increased sample size but at the expense of reduced false positive control leading to decreased precision.

In conclusion, we have developed novel ways of modeling the M-V data dependency and a new Bayesian statistical decision method for proteomics. Our Bayesian model is particularly robust for noisy data and shows improved performance over the state-of-the-art decision methods (3, 19, 20) and our M-V modeling improves the statistical decision methods.

## Results

### Bayesian data and decision model

Here we describe a Bayesian hierarchical model (Baldur) to analyze differences in peptide (or protein, PTM, etc., but for simplicity, we will use peptide from here on) abundances. This model will be applied to each peptide independently, and as such, we will describe it from the perspective of one single peptide. We assume the peptide’s data to be normally distributed with *C* different conditions (control/treatment, time points, etc.) each with *n*_*c*_ measurements. Then, we assume that the measurements within the *c*:th condition have a common mean *μ*_*c*_. We model each peptide’s data with a common standard deviation (sd) *σ* (unique to each peptide) and a measurement-specific uncertainty, *u*_*i*_ for the *i*:th observation, as a multiplicative factor that describes its change of variance from *σ* . Hence, the uncertainty is a correction for the unobserved measurement-specific variance around the mean. Further, we assume that all measurements and means in each condition are independent and therefore have zero covariance. We model the means with a group-level effect (21) and assume it is proportional to *σ* . To model this, we use the expanded non-centered parameterization which has been shown to increase sampling convergence and efficiency, and allows for increased model flexibility (21–27). This allows the model to adjust the posterior variance of each *μ*_*c*_ while still being constrained on *σ* and to shift the mean proportionally to *σ*. Finally, *σ* is assumed to be a gamma random variable with shape and rate parameterization, with hyperparameters estimated from the M-V trend. The data model is summarized in Equation 1.

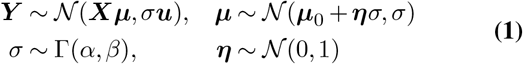

Here ***Y*** is a column vector of the N observation, ***μ*** is a column vector of the C means, ***X*** is an N-by-C design matrix, *σ*^2^ is the common variance, ***u*** is a column vector of the uncertainties, and ***η*** is a column vector (of length *C*) for group-level effects.

Baldur has two prior choices for the per condition ***μ***_0_, one empirical Bayes (EB) prior, and one weakly informative (WI) prior. The EB prior assumes a normal prior on ***μ***_0_ similar to the normal-normal compound model in (28, 29). The EB prior assumes that the mean of ***μ***_0_ is the sample mean with the variation set as twice the common variance normalized by the number of measurements (Equation 2). Here 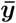 is a column vector of the sample means in the C conditions, and ***n***_*R*_ is a column vector of the reciprocal of the sample sizes in the C conditions. The WI prior uses a normal distribution with a substantial variance (Equation 3).

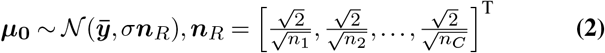

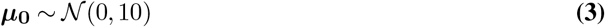

Next, we model our decision statistic ***D*** for comparisons of interest as a normal distribution with mean equal to a contrast of interest and variance equal to the common variance (*σ*^2^; Equation 4) normalized by the contrast weighted sample size, ***ξ***.

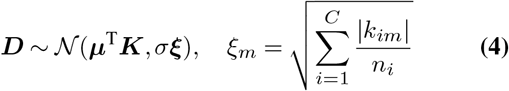

where ***K*** is a C by M contrast matrix (with M contrasts; Equation 5) of interest with the constraint that each column’s positive values sum to one and negative terms sum to minus one.

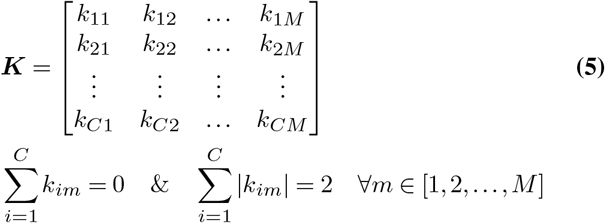

This allows for pairwise and non-pairwise comparisons, e.g., comparing the mean in one condition against the mean of two others. Here *σ* is invariant to the number of measurements, and ***ξ*** increases the confidence of a decision as the number of observations grows.

Finally, we estimate the probability of error by integrating the tails of ***D***. Let Φ be the cumulative density function of the standard normal distribution, ***μD*** the mean(s) of the posterior(s) of ***D, τ*** _***D***_ the reciprocal of the posterior sds (square root of the precision). Further, let the null hypothesis be that the difference in means is equal to *μ*_*h*_0 . Then the probability of error(s) is then defined according to Equation 6.

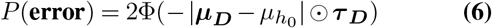

where ⊙ is the Hadamard product.

### Modeling the mean-variance trend

Here we will describe EB methods for estimating the hyperparameters of *σ* . Let 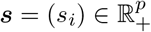 be a column vector of the sample standard deviations and 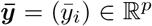 of the sample means in the *p* peptides of a dataset of interest. Here we use the sample standard deviation since the data model (Equation 1) assumes that each measurement has a unique variance.

#### Gamma regression for the mean-variance trend

The first inference model uses a gamma regression (GR) for estimating model hyperparameters. Let ***s*** be gamma-distributed and parameterized as described in Equation 7 (i.e., a GR with log-link function).

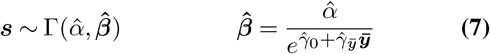

where Γ (., .) is the gamma distribution with shape, rate parameterization, and 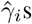 are the inferred regression parameters. Figure S1 shows the fitted GRs to the datasets investigated here (see the *Datasets* section in *Methods* for details). We found that the regression model describes the UPS-DS trend well. But, for the other datasets (in particular for the Human-DS) we observe that a single gamma regression model cannot capture the M-V distributions well.

We then define the uncertainty for some measurement *y*_*ij*_ as the expected sd (Equation 8).

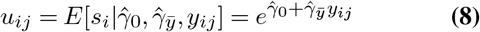

From Table 1, we see that the slopes and intercepts have similar values, except for the Human-DS. In addition, the shape parameter is slightly larger for this dataset compared to the other three. We then used the normalized root-mean-square error (Equation 9; NRMSE; (30)) to determine the goodness-of-fit.

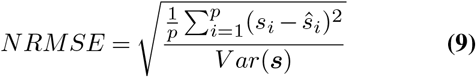

where *ŝ*_*i*_ is the predicted sd of the i:th peptide, and *V ar* (.) is the variance. We found that all datasets generated a similar error with the GR model.

**Table 1.**
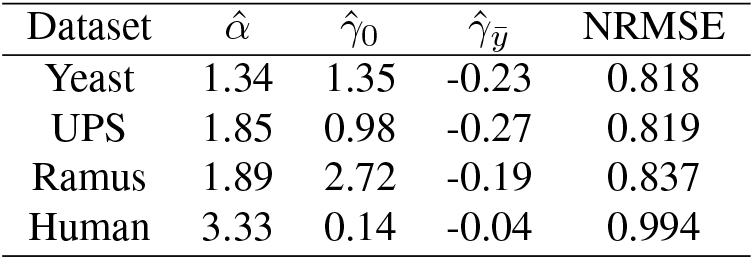
Table of inferred regression parameters and normalized root mean squared error for GR model.

#### Latent gamma mixture regression

To further increase the precision of the M-V trend modeling for each peptide, we propose a latent gamma mixture regression (LGMR) model. We assume that each peptide’s variance is a mixture of a common and a latent trend—allowing for localized estimates of the M-V trend parameters. The model starts with the same formulation as for the GR model (Equation 7 and 10) but with an exponentially modified normal distribution as a prior on *α* (Equation 11) and a folded normal hyperprior (Equation 12).

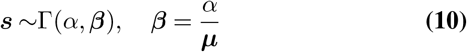

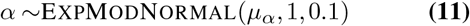

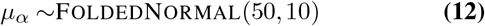

We then assume that the mean of ***s*** is a mixture of the two trends with one intercept and one slope each, and the *i*:th peptide has *θ*_*i*_ *∈* [0, 1] of the latent trend (Equation 13).

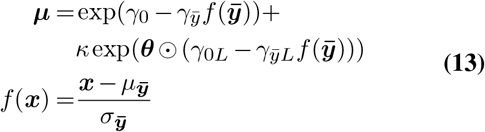

where *κ* is some small constant (here we will use 0.001) that defines the smallest possible contribution of the latent trend. We then choose a beta prior on *θ*_*i*_ (Equation 14) to enforce a peptide’s model towards either having a latent trend or not.

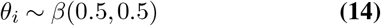

We set the slope to always be negative by limiting the slope coefficients to positive values 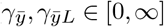; Equation 13). To this end, we used a half-normal prior on the slope coefficients (Equation 15-16).

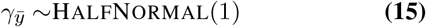

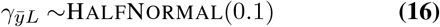

We then set priors on the intercepts (Equation 17-18). For the common intercept we used a standard normal distribution. For the latent intercept, we set a right-skew prior by setting the *α* parameter of the skew normal distribution to a large positive value. In addition, we set a fairly large variance by putting a large *ω* parameter to make the prior weaker. Finally, set the location parameter small positive for the latent intercept to force it larger and accommodate for the shrinkage by *κ*.

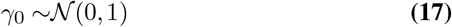

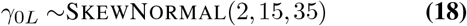

As for the GR model, we define the uncertainty for some observation *y*_*ij*_ as its expected sd (Equation 19).

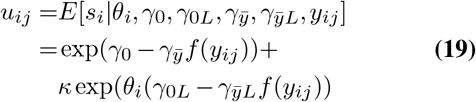

The inferred regression parameters of the LGMR model for the datasets investigated here are shown in Table 2 and visualized in Figure 1. We found that all datasets had unique regression patterns that resemble their corresponding M-V trend (Figure S1). From Table 2, we found that Ramus-DS and UPS-DS had similar *α* values, while the Yeast-DS value was smaller and Human-DS was significantly larger. We found that the Human-DS had the best fit followed by the Ramus-DS, UPS-DS, and Yeast-DS. Finally, compared to the GR model, we found that the LGMR model gave a better fit across all datasets (Table 1-2). Taken together, we have developed a new Bayesian regression model using a latent mixture that can describe well the local M-V trend of the datasets investigated here.

**Table 2.**
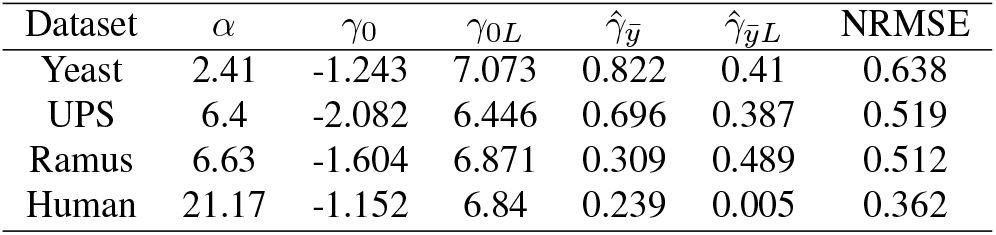
Table of inferred regression parameters and normalized root mean squared error for LGMR model. Numbers represents posterior means.

| Dataset | $\alpha$ | $\gamma_0$ | $\gamma_{0L}$ | $\hat{\gamma}_{\bar{y}}$ | $\hat{\gamma}_{\bar{y}L}$ | NRMSE |
| --- | --- | --- | --- | --- | --- | --- |
| Yeast | 2.41 | -1.243 | 7.073 | 0.822 | 0.41 | 0.638 |
| UPS | 6.4 | -2.082 | 6.446 | 0.696 | 0.387 | 0.519 |
| Ramus | 6.63 | -1.604 | 6.871 | 0.309 | 0.489 | 0.512 |
| Human | 21.17 | -1.152 | 6.84 | 0.239 | 0.005 | 0.362 |

**Fig. 1.**
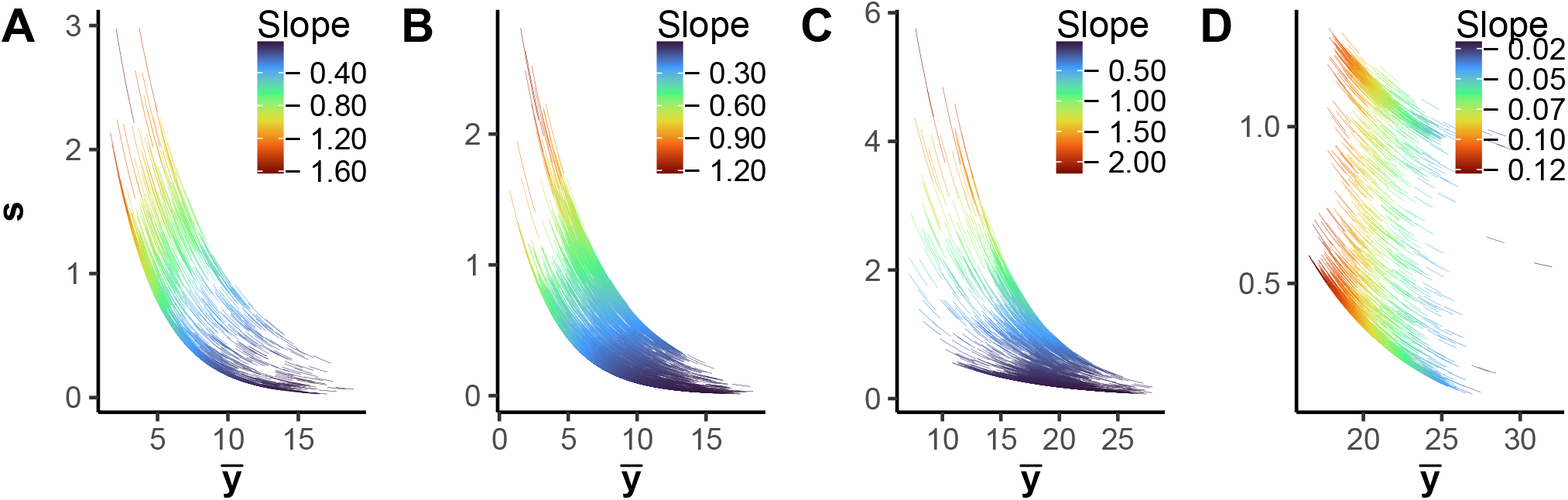
The mean-variance trend in the yeast-, UPS, Ramus-, and Human-DS (**A**-**D**, respectively) with the locally estimated gamma regression. X-axis shows the sample mean, Y-axis shows the sample standard deviation, each line indicates the M-V trend of a corresponding peptide, and the color indicates the derivative at the peptide’s mean.

### Algorithmic description

The procedure for implementing the Bayesian decision is described in algorithm 1. The user needs to input their data, a design matrix, a contrast matrix, a choice of regression model to use, LGMR or GR, and finally a choice of prior, EB or WI. Baldur then fits the regression model and uses it to infer uncertainties as well as priors parameters on *σ*. It then runs the decision model on each peptide separately and produce a summary statistic of the fit (this allows for a highly efficient parallel computation setup). In particular, Baldur returns the mean, median, a 95 % credibility interval, the R-hat, and the efficient sample size for parameters of interest.

#### Algorithm 1

description of Baldur

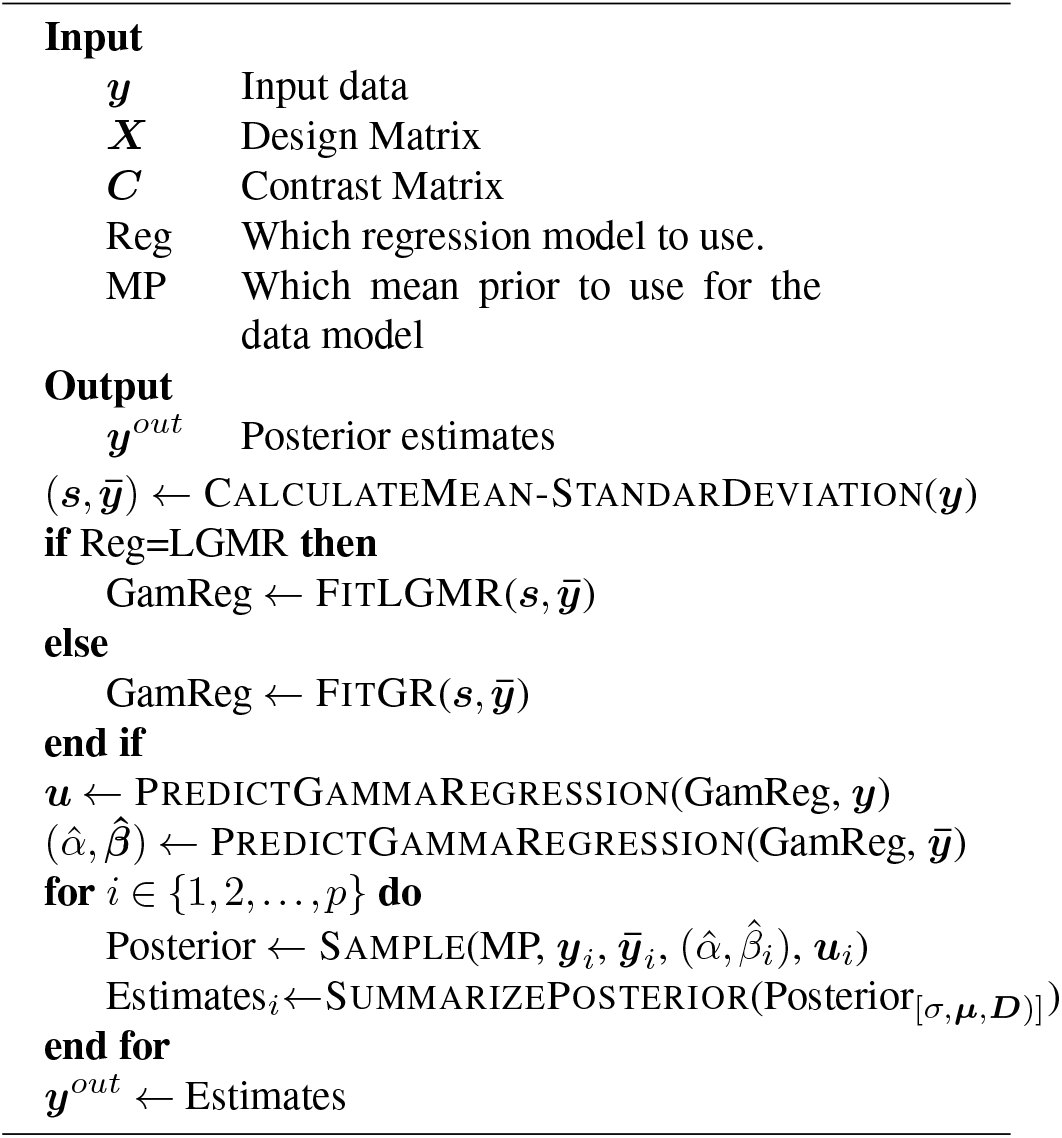

### Performance evaluation

#### Receiver Operator Characteristics

To evaluate the performance of the models presented here, we generated receiver operator characteristic (ROC) curves of the four benchmark datasets. We evaluated the following methods: Baldur with both priors and regressions, Limma-Trend, and t-test since they are generally used in recent studies (3, 14, 19, 20, 31– 34). For Limma-Trend and t-test, we applied false discovery rate correction using the method described in (35). For the Yeast-DS (Figure 2), we observed an increased performance using Baldur. In particular, all Baldur-based models improved on Limma-Trend and t-test. In addition, the LGMR model for parameter inference slightly improved on the GR model, and the weakly informative (WI) prior for the mean had a marginally better performance than the empirical Bayes (EB) prior.

**Fig. 2.**
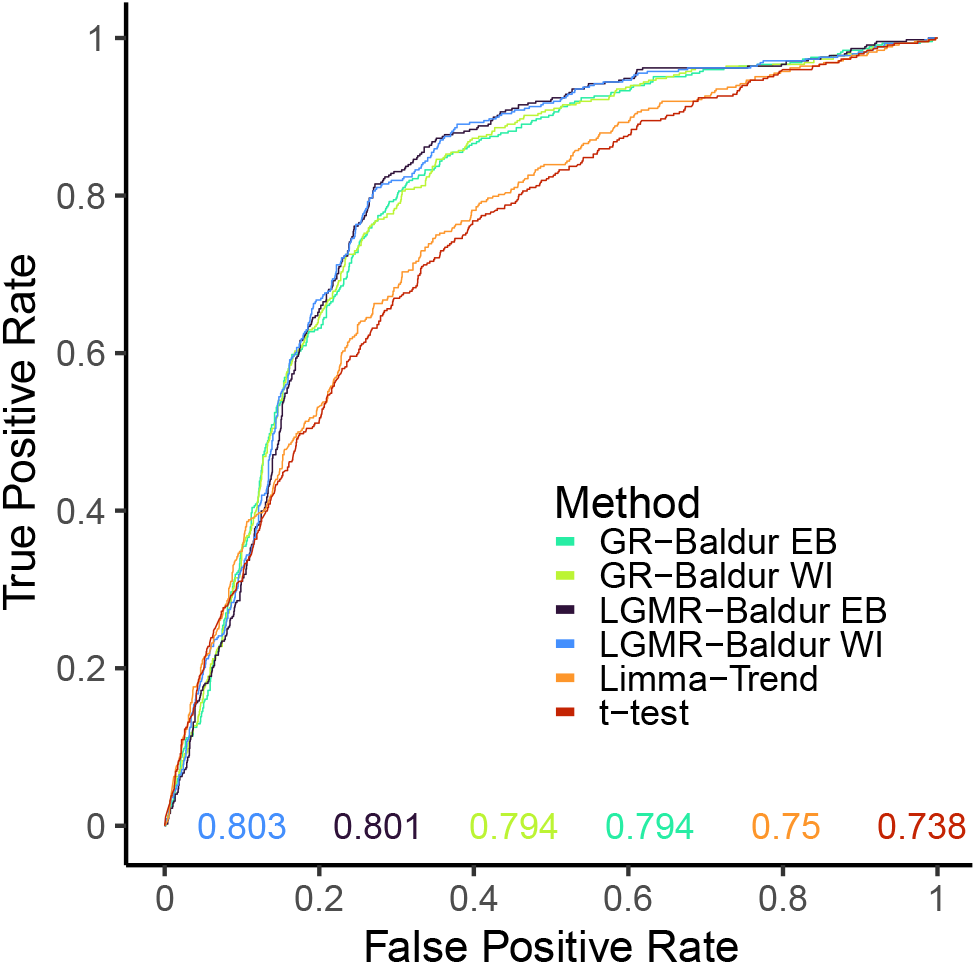
Receiver operator characteristic (ROC) curves for the Yeast-DS. The different colors indicate the evaluated models; LGMR-Baldur is the Baldur method with the LGMR model (Equation 10-18) to estimate *σ* priors (Equation 4) and uncertainty (Equation 19), GR-Baldur is using Equation 7 to estimate *σ* priors and uncertainties (Equation 8), EB and WI indicates the empirical Bayes prior (Equation 2) or the weakly informative prior (Equation 3) on the mean, Limma-Trend as described in Methods, and t-test used with pooled variance. The X-axis shows the false positive rate (Equation 22), and the Y-axis shows the true positive rate (Equation 23). The colored numbers on the bottom of the plots show the area under the curve with colors corresponding to the different models.

For the UPS-DS we show the ROC curves for both pairwise and non-pairwise comparisons (Figure S2**A**). While all methods generally performed well on the pairwise comparisons, Baldur with the LGMR model has a notably better performance in all comparisons, Baldur with the GR model is second, followed by Limma-Trend and t-test. In addition, we found that Baldur-based models had a similar area under the ROC curve (auROC) across all pairwise comparisons, while Limma-Trend and t-test had a larger spread in performance for the different comparisons. In addition to pairwise comparisons, we also studied non-pairwise design for the UPS-DS to evaluate the performance of the statistical decision at small and intermediate average log fold changes of the spike-in peptides. We found that all methods showed similar performance for the larger fold-changes, but for the very small fold-change of *fmol50 vs fmol25 and fmol100* (i.e., the fold-change of true positives are 0.8) we found that LGMR-Baldur methods outperformed GR-Baldur which had a slightly better or equal performance to Limma-Trend. In addition, the EB prior showed slightly better performance than the WI prior. For the Human-DS (Figure S2**B**), we found that all methods had similar performance across all comparisons except one (1:12 vs 1:6), where LGMR models had slightly better performance.

For the Ramus-DS (Figure 3), we identify a wide range in performance for different comparisons ranging from easy (e.g., 5 vs 50) to hard (e.g., 0.125 vs 0.5). Still, we found that both GR and, in particular, LGMR Baldur models can improve performance significantly when the spike-in quantities are in low concentrations and involve smaller fold-changes between conditions. In particular, we see performance improvements for comparisons where the spike-in concentration is lower than 12.5 fmol. Conversely, we find comparable performance for all methods when the fold changes are substantial or when the spike-in concentrations are ample. Still, it is evident that Baldur with the LGRM model’s performance is highest in all 36 comparisons, while GR-based inference is second.

**Fig. 3.**
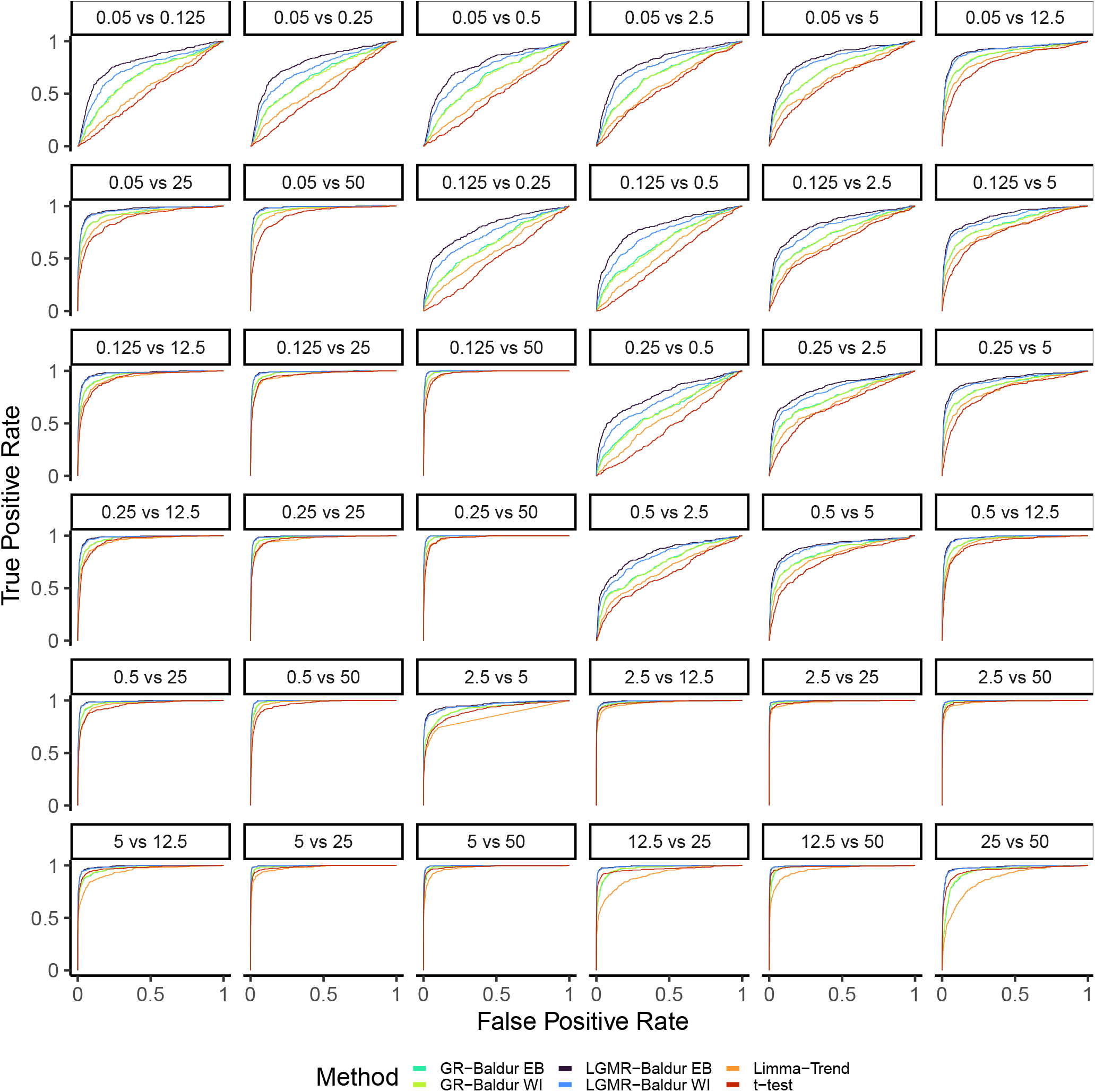
Receiver operator characteristic curves for the comparisons in the Ramus-DS. Facet titles indicate what comparison is being evaluated together with the fmol spike-in concentration of the UPS1 of the two conditions. Color, axis, numbers, points, and decision as described in Figure 2.

Summarising, Baldur has drastic performance improvement in noisier conditions of proteomics measurements, with the LGMR best-performing alternative and GR second. On the other hand, for less noisy (easier) datasets (or comparisons), Baldur is marginally better than Limma-Trend or t-test, which already have large auROC and therefore have little room for improvement. In addition, we observed that the LGMR model consistently improves Baldur’s performance over GR model inference. Finally, we observed that Baldur has dependable performance across all datasets, while Limma-Trend and t-test can show considerable variation in performance between comparisons.

#### Decision patterns

While ROC curves can produce a summary statistic over a classifier’s support—it is inadequate for a deeper understanding of specific statistical thresholds typically used in the significance analysis of proteomics data. Thus, we investigated Matthews correlation coefficient (MCC; Equation 25; (36, 37)) as a point estimates of statistical decisions over a range of significance levels. The motivation for using MCC is that all datasets are heavily unbalanced between true positives and true negatives (Table 3), and MCC has, arguably (38, 39), good properties for unbalanced data. For Yeast-DS (Figure S3), we found that LGMR does performed slightly better than GR and the EB is better paired with LGRM-based inference while WI works better with GR-based parameter estimation. In addition, we found that Baldur methods always have a larger MCC compared to Limma-Trend and t-test. Finally, we found that the Baldur methods make an optimal decision around 1 to 5 % while Limma-Trend and t-test peak very close to 0 and subsequently decay with the significance level.

**Table 3.** Properties of the datasets investigated here. TP (True Positives) is the number of spike-in examples, TN (True Negatives) the size of the background, Total is the sum of the two, Conditions is the number of different spike-in confrontations, and Replicates is the number of replicates per condition.

| Dataset | TP | TN | Total | Conditions | Replicates |
| --- | --- | --- | --- | --- | --- |
| Yeast | 448 | 1787 | 2235 | 2 | 3 |
| UPS | 392 | 10207 | 10599 | 3 | 4 |
| Ramus | 292 | 4654 | 4946 | 9 | 3 |
| Human | 404 | 2902 | 3306 | 3 | 23 |

Similarly to the ROC curves, the Ramus-DS showed a wide range of performance for the MCC (Figure S4) between different comparisons. Still, Baldur using GR and LGRM inference tended to produce better-performing decisions that generally decay slower with the significance level compared to Limma-Trend and t-test that often peak at very small significance levels. Importantly, while some ROC curves (Figure 3) suggested similar performance, the MCC showed substantial changes at typical significance levels. In particular, we see that Baldur-based models showed better performance when compared to Limma-Trend and t-test in the 5 vs 50 comparisons; all methods showed good ROC performance, while Limma-Trend and t-test have significantly decreased MCC.

Finally, we see that the Bayesian decision tends to have robust performance while t-test and Limma-Trend have unexpected drops in performance, in particular at lower significance levels.

For the UPS-DS (Figure S5), we again found that Baldur-based decisions tend to outperform Limma-Trend and t-test. In all pairwise comparisons, we find that the LGRM model tends to produce the best decisions. In particular, we find that the WI prior slightly increases performance at the comparison *fmol25 vs fmol100* that has the larger fold-change. We find that while the GR method is slightly worse than Limma-Trend for the *fmol50 vs fmol100* and *fmol25 vs fmol50* comparisons it shows better performance in the *fmol25 vs fmol100* comparison. For the non-pairwise contrasts, we found that the LGMR and GR models performed better at typical significance levels. For the most challenging contrast (*fmol50 vs fmol25 and fmol100*) we find a significant drop in Baldur performance, while Limma-Trend does not produce any decision until a large significance level (greater than 0.19).

For the Human-DS (Figure S6), we observed that LGMR improves the decision over Limma-Trend and t-test in all three comparisons, while the GR model has better performance in two comparisons. Surprisingly, Baldur performs considerably better at the largest fold change where Limma-Trend and t-test rapidly drop in performance. In addition, we observe that both priors behave (almost) identically in all three comparisons, likely due to the large number of replicates in this dataset.

In conclusion, from the MCC evaluation, we found that Baldur tends to make more balanced decisions around traditional significance levels. In addition, we found that Baldur retains a more balanced decision over a wide range of decisions compared to Limma-Trend and t-test and is particularly good in comparisons where all models have their lowest performance (i.e., has the best worst-case performance). Finally, we found that both EB and WI generally perform similarly from a balanced decision perspective. Still, the LGRM model shows the highest performance across all datasets and comparisons independent of prior choice.

Next, we analyzed the true-positive rate (TPR), the false-positive rate (FPR), and precision as a function of the significance level. For the Yeast-DS (Figure 4), we see an improvement, foremost in controlling the FPR, while still being competitive in TPR. While all models produce large FPR, Baldur shows robust control at small significance levels and slowly increases in FPR. On the other hand, Limma-Trend and t-test rapidly pick up false positives at lower significance levels. In particular, we observe that LGMR model controls the FPR well for both priors, while the GR inference with the EB prior tends to produce the largest TPR and FPR of the Baldur methods. Importantly, the control of the FPR leads to increased precision of the Baldur methods, all improving over Limma-Trend and t-test.

**Fig. 4.**
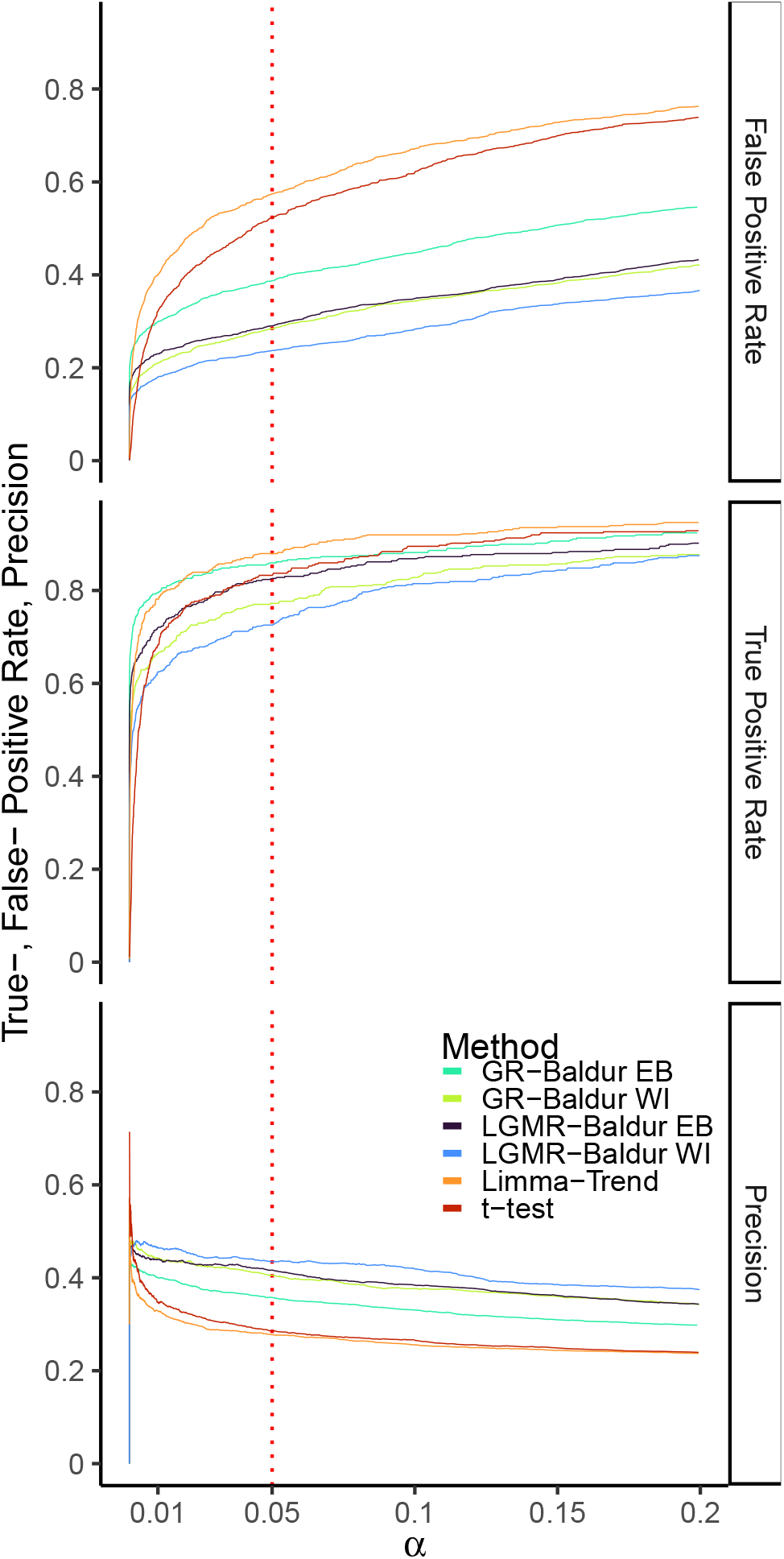
Performance metrics of the Yeast-DS plotted against the significance level (*α*). Y-axis shows the metric value (as indicated by facet titles), X-axis shows the significance level, and colors as according to Figure 2.

For the Ramus-DS (Figure S7-S8), we again observed that Baldur models tend to control the FPR to a much higher degree while having similar or even better TPR. In particular, we observed that the LGMR-EB setup vastly outperforms Limma-Trend in reducing FPR while attaining marginal improvement in TPR. We observe that the improvements in Baldur methods come from better control of the FPR rather than an increase in TPR. In addition, Baldur controls the FPR robustly in all comparisons, while Limma-Trend and t-test have fluctuating FPR.

Next we investigated the performance of the pairwise statistical decisions in the UPS-DS (Figure S9) which has the highest imbalance of the datasets investigated here (Table 3). We found that the LMGR had the best performance in terms of TPR, FPR control, and precision in all three comparisons. While the GR model had lower TPR, its FPR control still produced the second best precision in all comparisons. For the non-pairwise contrasts (Figure S10), we observed a similar pattern in performance where LGMR had the best performance, GR had a lower TPR while still maintaining among the best precision; similarly, Limma-Trend produced a significantly higher FPR. The third comparison at low spike-in log fold change (-0.32) all methods had very low TPR. Limma-Trend made no decision until a large significance level of roughly 0.18, while LGMR-EB managed to produce the largest TPR among all methods.

For the Human-DS (Figure S11), we saw that Baldur methods have outstanding precision over Limma-Trend and t-test due to the strong FPR control in this dataset. Further, all models can get a high TPR, except for GR in *1:12 vs 1:6*. Still, and particularly for the Human-DS, we observed that, in the worst-case scenario, Baldur-based models make an improved decision by better controlling the FPR. Since these datasets have large TN, this translates to increased precision for all Baldur methods over Limma-Trend, with the WI prior slightly outperforming the EB.

Concludingly we observed that investigating performance metrics as a function of significance level can elucidate model-specific behavior. In particular, Baldur’s methods tend to control the FPR much better than t-test and Limma-Trend while generally having comparable TPR. This lead to the best preclusion in all dataset and comparison for Baldur. Concurrently, we found that this led to an improvement in a balanced decision as measured by the MCC, indicating that Baldur’s methods tend to make more balanced decisions. This becomes obvious when examining the distribution of FP and TP at the 5 % significance level for all four datasets (Figure 5 and Figure S12-S14) where Baldur based methods tend to have similar TP but far lower FP. The exemption is for a few comparisons in the Ramus-DS and in the UPS-DS where the LGMR routine picks up more TPs and fewer FPs.

**Fig. 5.**
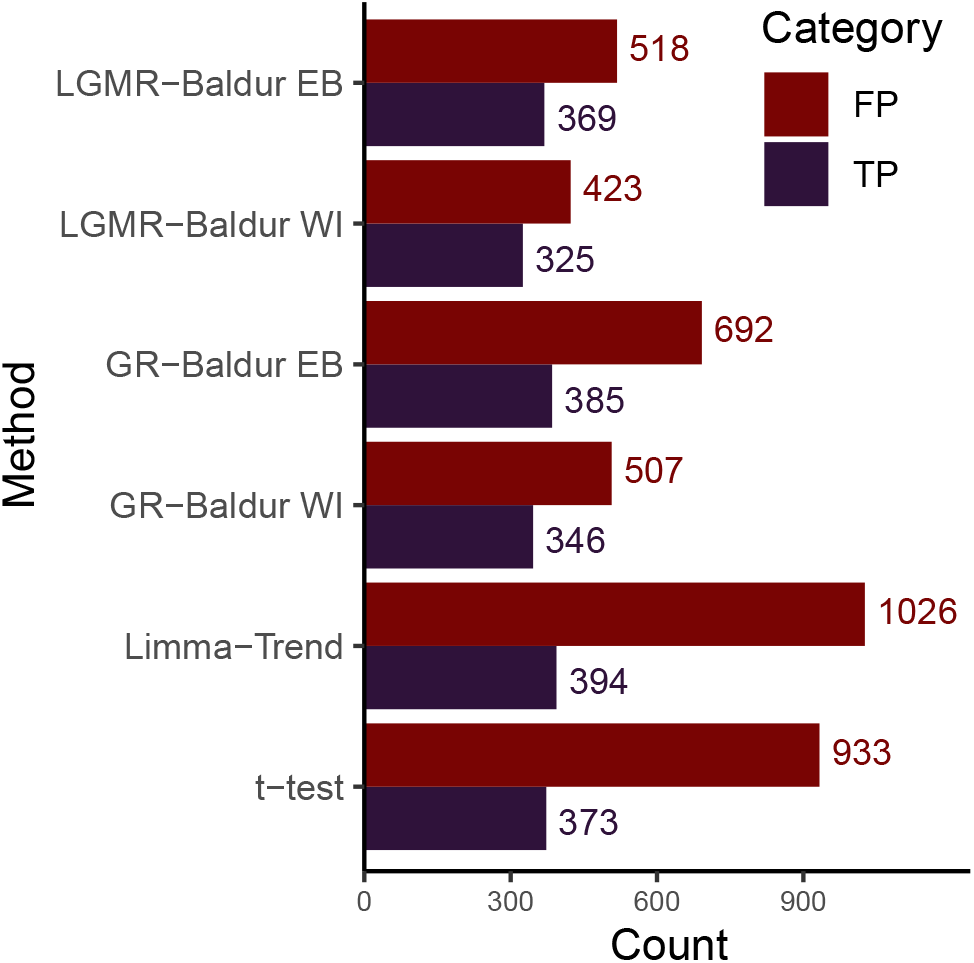
Decision of False Positives and True Positives of at 5 % significance level for Yeast-DS and the methods evaluated here.

#### Empirical power analysis

As a final analysis, we investigated the performance of the methods presented here as a function of replicates. The large number of replicates in the Human-DS (Table 3) allowed for analyzing how the number of replicates affects the statistical decision. To this end, we varied the number of replicates from 3 to 21 within each condition by producing 24 randomly selected combinations for each replicate size. We then investigated the TP and FP at a 5 % significance level for each replication size and combination. For the first comparison, we found that all methods except t-test could identify most TPs at three replicates, and all methods could identify all TPs at six replicates (Figure 6). Surprisingly, as the number of replicates increased, both t-test and Limma-Trend increased in the number of FPs. On the other hand, Baldur methods remained steadily at the same number of FPs for the entire range of replicates; this holds for all three comparisons in the dataset. For the second comparison, we again see that t-test and Limma-Trend increase in FPs as the number of replicates increase but flatten around 18 to 21 replicates. In addition, we see that GR with both priors have subpar TP detection compared to the others that identified nearly all TPs at around 18 replicates. Finally, we observed that in the third comparison LGMR and Limma-Trend could identify more TPs than other methods at three replicates. At nine replicates, we observed that all methods identified most of the TPs and that there was no change in FPs as replicates increased. Taken together, Baldur’s decision benefits from increasing the number of replicates. On the other hand, increasing the number of replicates can reduce the performance of t-test and Limma-Trend by reducing control of the FPR.

**Fig. 6.**
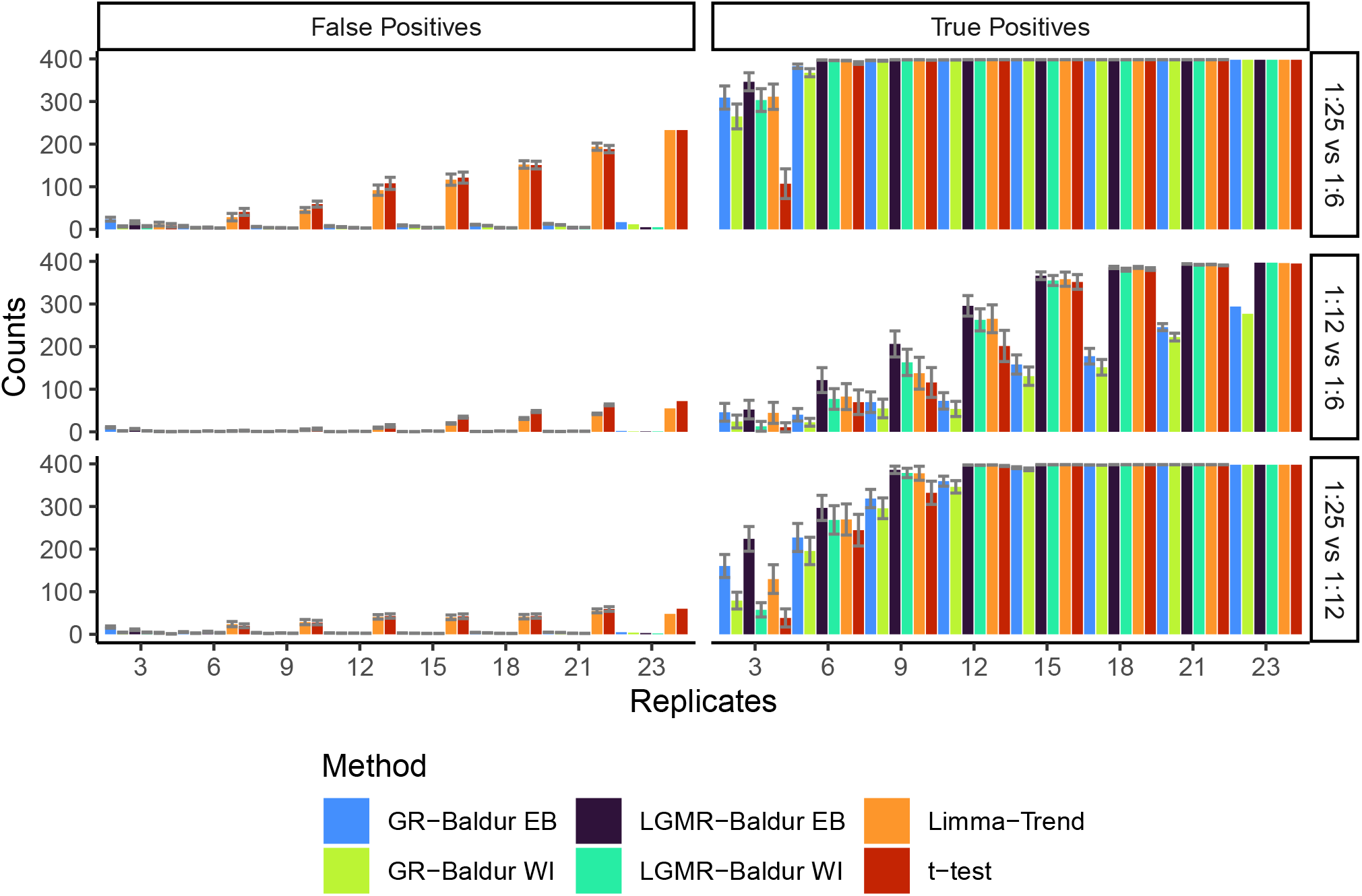
Baldur increases in power with increasing number of replicates without the expense of increased false positives in the Human-DS. Bars show the average number of false- or true-positives (indicated by facet titles) at a 5 % significance level of n=24 independent runs with each run having a unique subset of replicates. Error bars shows the mean *±* the standard error of the mean, 24 replicates are all replicates in the data and therefore a point estimate. X-axis indicates the number of replicates used. Colors as in Figure 2.

## Discussion

While increasing TPR is a driver for proteomics discovery, controlling the FPR is critical for precise high-throughput functional analysis of proteomes and reactomes and enables a better downstream systems biology analysis. In addition, increased FPR control will be directly associated with improved reproducibility of a study. Here we demonstrate that Baldur with the LGMR model has the best performance across all comparisons in all four datasets investigated here despite only one of them exhibiting a strong mixture of distributions (Figure 1 and Figure S1).

We compared two different priors for the mean of each condition, a weakly informative (WI; Equation 3) and a data-driven Empirical Bayes (EB; Equation 2). In general, we found comparable performance, but the WI prior tends to produce a smaller TPR while having a higher precision. On the other hand, the EB prior tends to produce higher TPR but with lower precision. Compared to t-test and Limma-Trend, the EB prior has a comparable TPR but reduced FPR, while the WI has lower TPR but even smaller FPR. Further, we find that the inference of constants and hyperparameters for Baldur with the latent gamma mixture regression (LGMR) model developed here outperforms gamma regression (GR) in all comparisons. Likely, this is due to the improved M-V trend modeling of the LGMR model being able to infer the local M-V trend for each peptide in the datasets.

While the LGMR model developed here (Equation 11-18) fits and describes well all datasets analyzed in this study (Table 3, Figure 1 and Figure S1), there is no guarantee that it will fit any input dataset produced by the vast number of spectral quantification methods. Still, in the four datasets investigated here (Figure S1) we found that the LGMR model described the M-V trends well (Figure 1 and Table 2). In addition, using Baldur with a GR model tends to outperform t-test and Limma-Trend and pose as a valid option for the LGRM at a lower computational cost.

Lastly, we believe that Baldur’s robust control of false alarms will be highly effective for analyzing PTM and peptidomics datasets susceptible to have larger sample variations. Here we observed this in Yeast-DS (PTM dataset) and Ramus-DS for small spike-in concentrations and in conditions with small fold-changes between TPs.

## Conclusions

We have developed a new data-driven methodology to analyze label-free proteomics data. The proposed methods build on modeling the variance component using a gamma random variable. We introduce a new Bayesian decision method (Baldur) which improves performance over previously published proteomics data analysis methods by controlling the false alarm. Further, we proposed a novel regression model to characterize the mean-variance dependency observed in the proteomics data. When experimental data is noisy, as is common in PTM and peptidomics analyses, we expect that Baldur with the latent gamma mixture regression model will have best performance. Future work should focus on approximative Bayesian methods to speed up computational analysis for big datasets.

## Methods

### Datasets

We used four previously published (14, 19, 20) label-free spike-in benchmark datasets to evaluate model behavior and performance. Two benchmark data sets (19) consists of LC-MS/MS: one on total proteomics with the Universal Proteomics Standard Set 1 (UPS1) spiked in at 1:2:4 times concentration in a Chlamydomonas background, and the other is a PTM quantification dataset using a reversibly oxidized cysteine enrichment protocol with *Saccharomyces cerevisiae* spiked in at 1:2 concentrations to a Chlamydomonas background (see original paper for details). We referred to these datasets as UPS-DS and Yeast-DS, respectively. The UPS-DS has four replicates per spike-in (condition), and Yeast-DS has three. In addition, we used the data published by (14, Ramus-DS) that has UPS1 spiked in at nine different concentrations to a yeast background with three replicates in each condition. Finally, we investigated a recently published dataset (20) where *Escherichia coli* proteome was spiked-in at 6:12:25 concentrations to a background of a heterogeneous human tumor population. Table 3 summarizes the properties of these datasets.

### Data pre-processing

For the Yeast-, UPS-, and Ramus-DS, normalization was done by calculating the scale factors described in (40), and dividing each sample accordingly. That is, let *y*_*ij*_ be the measurement of the *i*:th peptide in the *j*:th sample. The normalization constant *s*_*j*_ for the *j*:th sample is then given by equation 20

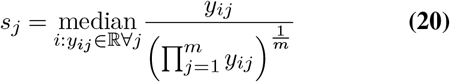

The normalized data 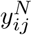 was then given by equation 21.

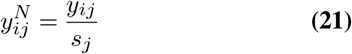

Where *s*_*j*_ were only calculated using rows without missing values, and after normalization, the data was log_2_ transformed. Lastly, Yeast-, UPS-, and Ramus-DS were imputed using missForest (41). For the Human-DS, columns corresponding to conditions with no spike-in was discarded, then rows with missing values were filtered out prior to normalization, to be in accordance with the arguments presented in the original study (20). For the empirical power analysis, the columns were subset from the normalized data.

### Model inference

For inference of the latent gamma mixture regression model, we set *κ* = 0.001. Then, we used RStan’s (42) No-U-Turn Sampler (NUTS) using five chains each with 1500 warm-up draws, 4500 post-warm-up samples (iter set to 6000), adapt_delta set to 0.9 for everything except the empirical power analysis. For the empirical power analysis, we ran 20 chains with 1000 warn-ups draws and 1500 post-warm-up samples. For the calculation of the normalized root-mean-square error, Equation 9 was calculated at each iteration. The sample mean from the posterior was then used as a point estimator and presented in Table 2. Inference of the posterior distribution for the data and decision model was done with RStan’s NUTS using four chains each drawing 1000 burn-ins and 1000 samples per peptide. The parameters for the gamma regression was estimated using R’s (43, Version 4.2.0) function **glm** (44), and the shape parameter was estimated using the R package stats’s **summary.glm** function. For integrating ***D*** (Equation 4), we used the normal cumulative distribution function as implemented in R’s **pnorm** function using the sample mean and sample standard deviation from the posterior draws.

### Running limma and t-test

For running Limma-Trend, we first ran **lmFit**, then **contrast.fit**, and **eBayes** was ran with Robust and trend set to TRUE. The p-values were then extracted with **topTable** with adjust.method set to “fdr” and number set to Inf. For t-test, we used R’s **t.test** function with var.equal = TRUE, and then ran **p.adjust** with method set to “fdr” on the p-values (within each comparisons).

### Performance metrics

For performance, all decisions were calculated over the closed set [−0.1, 1] so the ROC curves always starts at the origin. True positive rate was calculated according to equation 23, false positive rate according to equation 22, precision according to equation 24, and Mathews correlation coefficient according to equation 25.

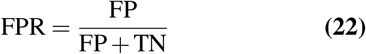

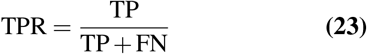

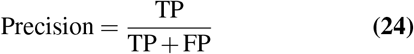

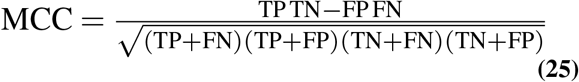

## ACKNOWLEDGEMENTS

We like to acknowledge Hicks lab at the University of North Carolina at Chapel Hill, and in particular former PhD student Dr. Evan McConnell, for providing and describing the Yeast-DS, UPS-DS, and Ramus-DS.

## Availability of data and materials

Baldur is available through GitHub (https://github.com/PhilipBerg/baldur) and through The Comprehensive R Archive Network (**install.package(“baldur”)**). Yeast-DS, UPS-DS, and Ramus-DS datasets are available through the figshare repository (https://figshare.com/s/28e837bfe865e8f13479); Yeast-DS as well as UPS-DS are included in the Baldur package. The Human-DS is available through the original publication (20). The R code used to produce this paper is available through GitHub (https://github.com/PhilipBerg/baldur_code).

## Authors’ contributions

PB and GP designed the statistical methods and conceived the study. PB designed the R package, performed the analysis, and wrote the corresponding R code. PB wrote the paper. GP and PB edited the final manuscript.

## Dedication

The authors would like to dedicate this paper to the memory of Professor Sorina Popescu.

## Supplementary Figures

**Fig. S1.**
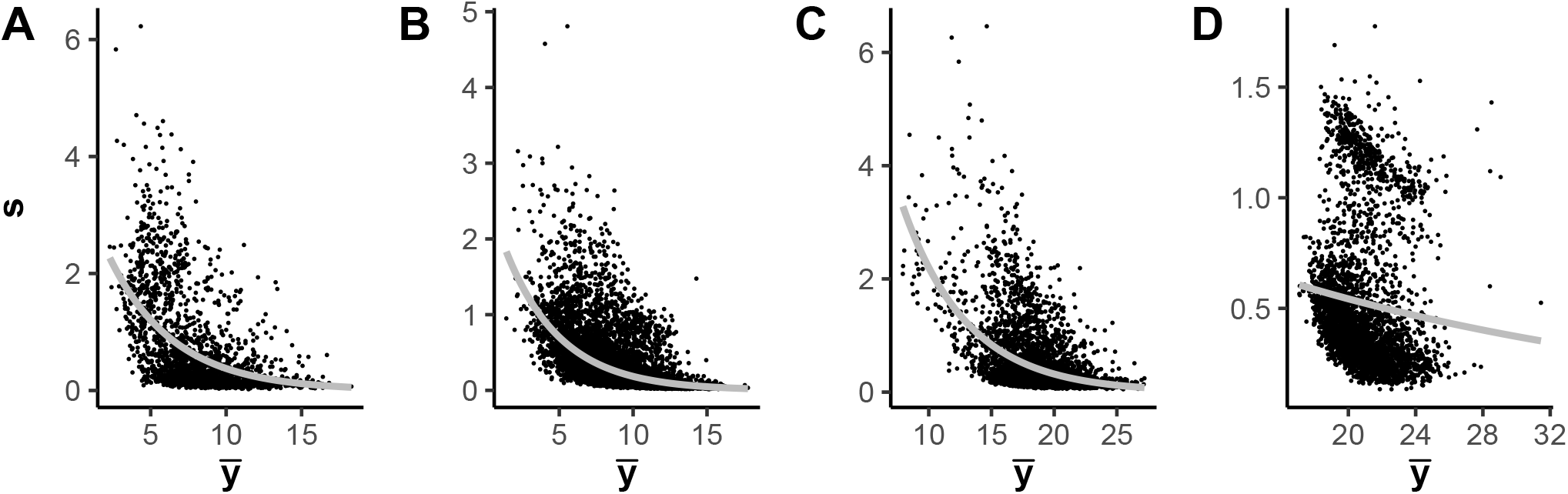
Mean-variance trend in the Yeast-, UPS-, Ramus-, and Human-DS (**A**-**D**, respectively) with the estimated gamma regression. X-axis shows the sample mean, and Y-axis shows the sample standard deviation, and the gray line represents the estimated gamma regression model.

**Fig. S2.**
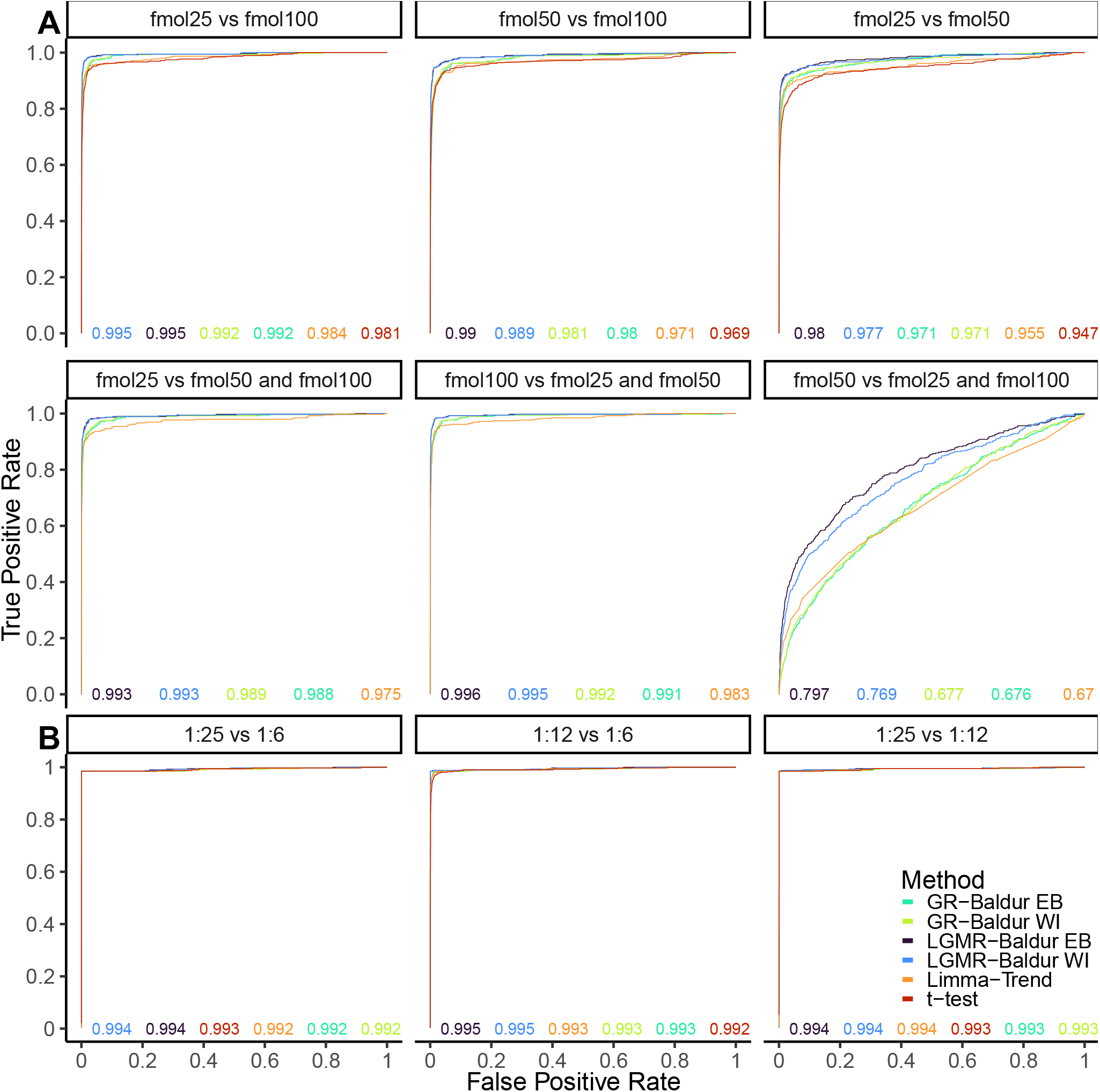
Receiver operator characteristic curves for the three comparisons in the UPS-DS (**A**) and Human-DS (**B**). Facet titles indicate what comparison the plot graphs. For the UPS-DS, titles show the spike-in concentrations of UPS1 (25-, 50-, or 100-fmol; 1:2:4), and, for the Human-DS, they show the human to *e. coli* peptide ratio (i.e., human:*e. coli*). “and” indicates the mean of the two conditions (e.g., contrast vector [-1 0.5 0.5]^T^). Color, axis, and numbers, as described in Figure 2.

**Fig. S3.**
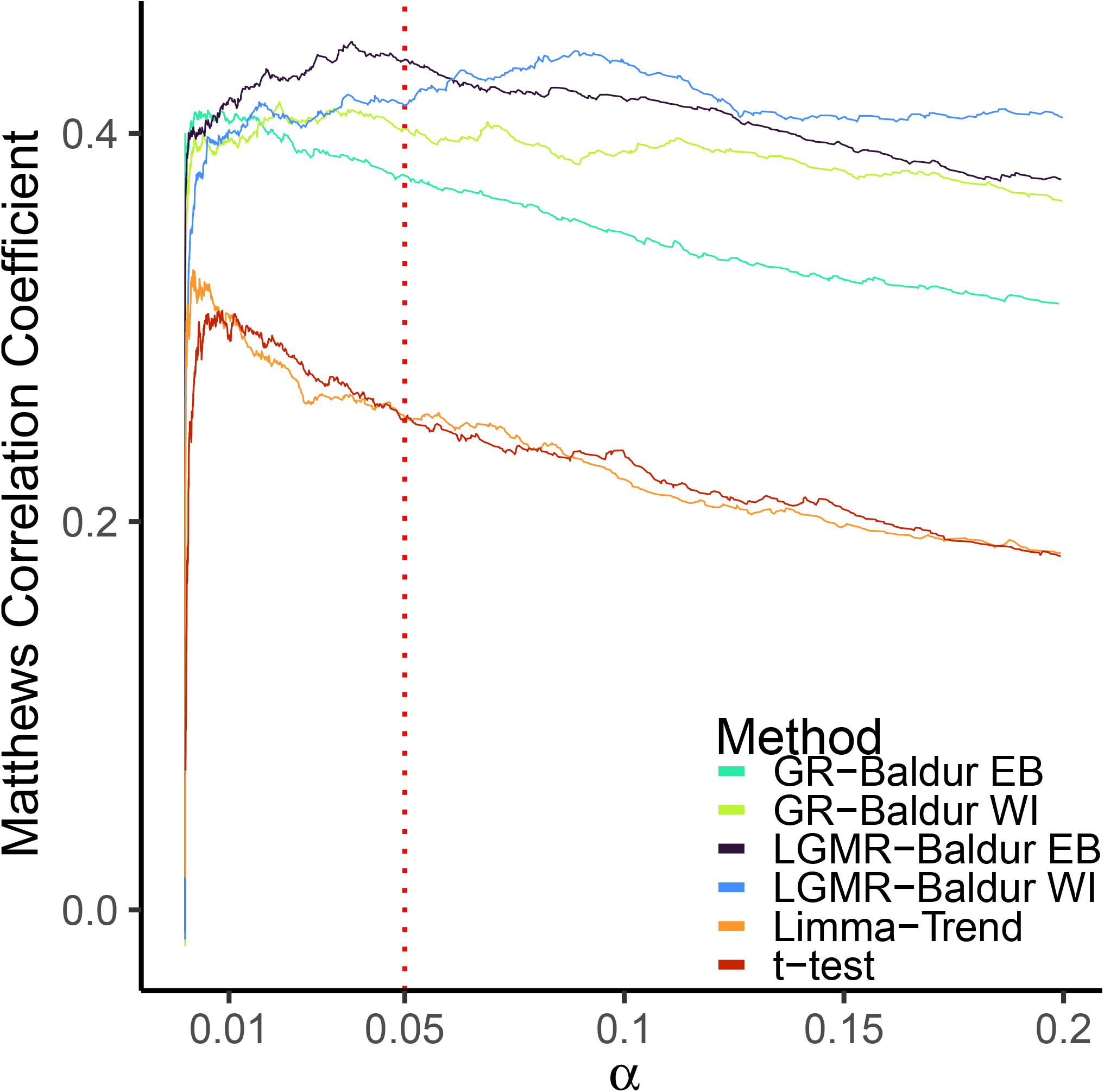
Mathews correlation coefficient of the Yeast-DS plotted against the significance level (*α*). Y-axis shows the Mathews correlation coefficient, X-axis shows the significance level for the different comparisons (as indicated by the Y-axis facet titles), and colors as according to Figure 2.

**Fig. S4.**
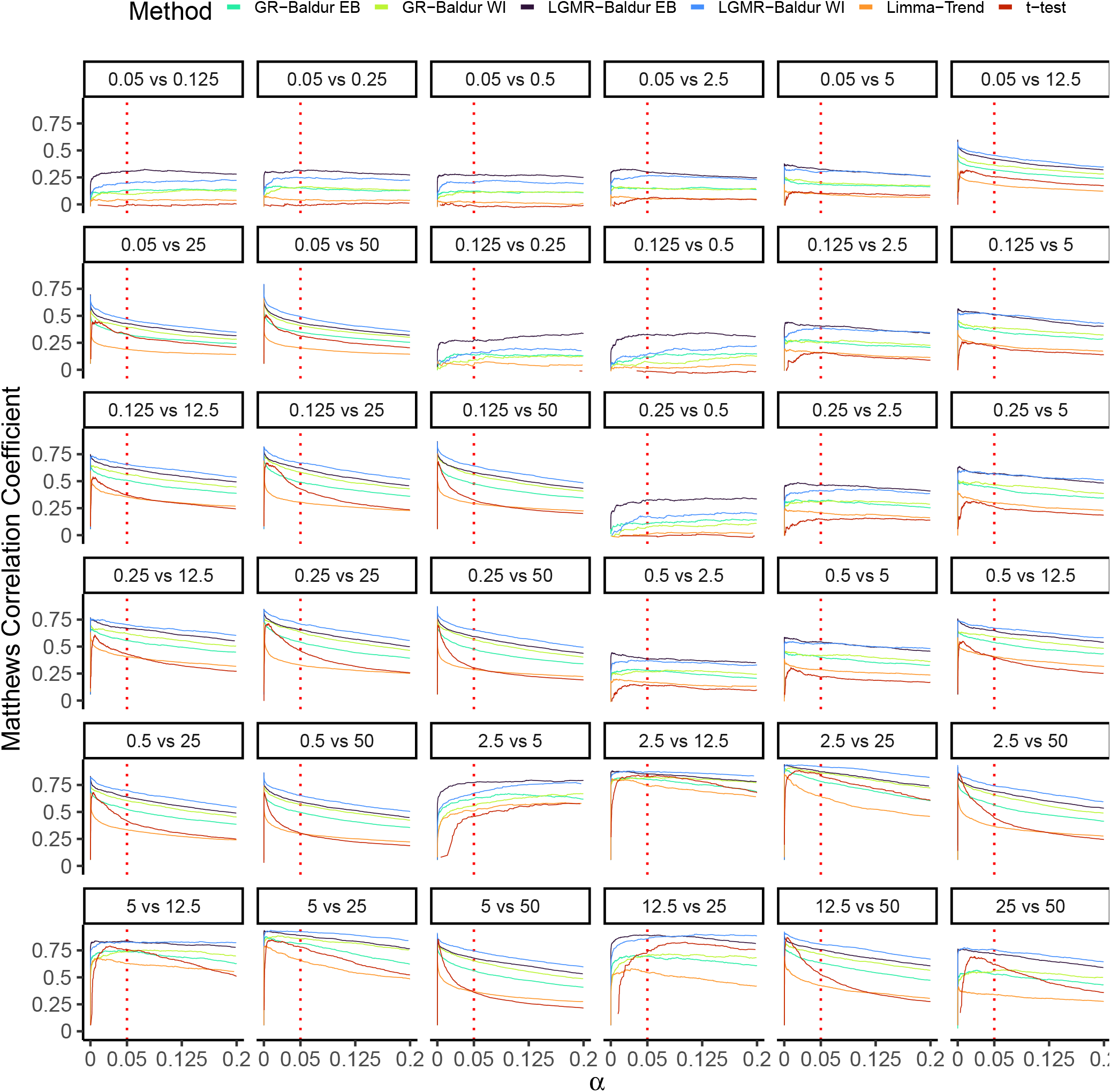
Mathews correlation coefficient of the Ramus-DS plotted against the significance level (*α*). Y-axis shows the Mathews correlation coefficient, X-axis shows the significance level for the different comparisons (as indicated by the Y-axis facet titles), and colors as according to Figure 2.

**Fig. S5.**
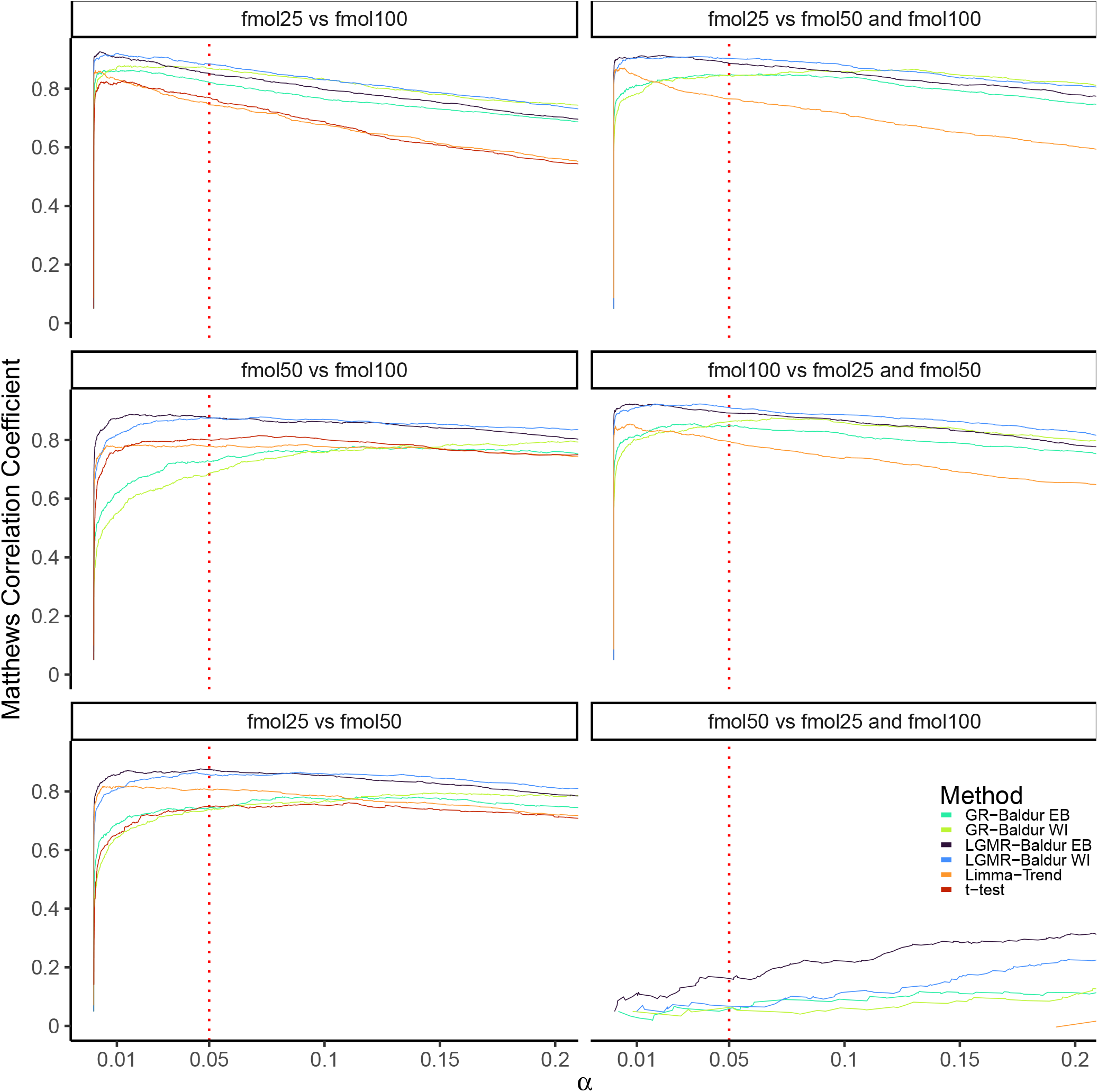
Mathews correlation coefficient of the UPS-DS plotted against the significance level (*α*). Y-axis shows the Mathews correlation coefficient, X-axis shows the significance level for the different comparisons (as indicated by the Y-axis facet titles), and colors as according to Figure 2.

**Fig. S6.**
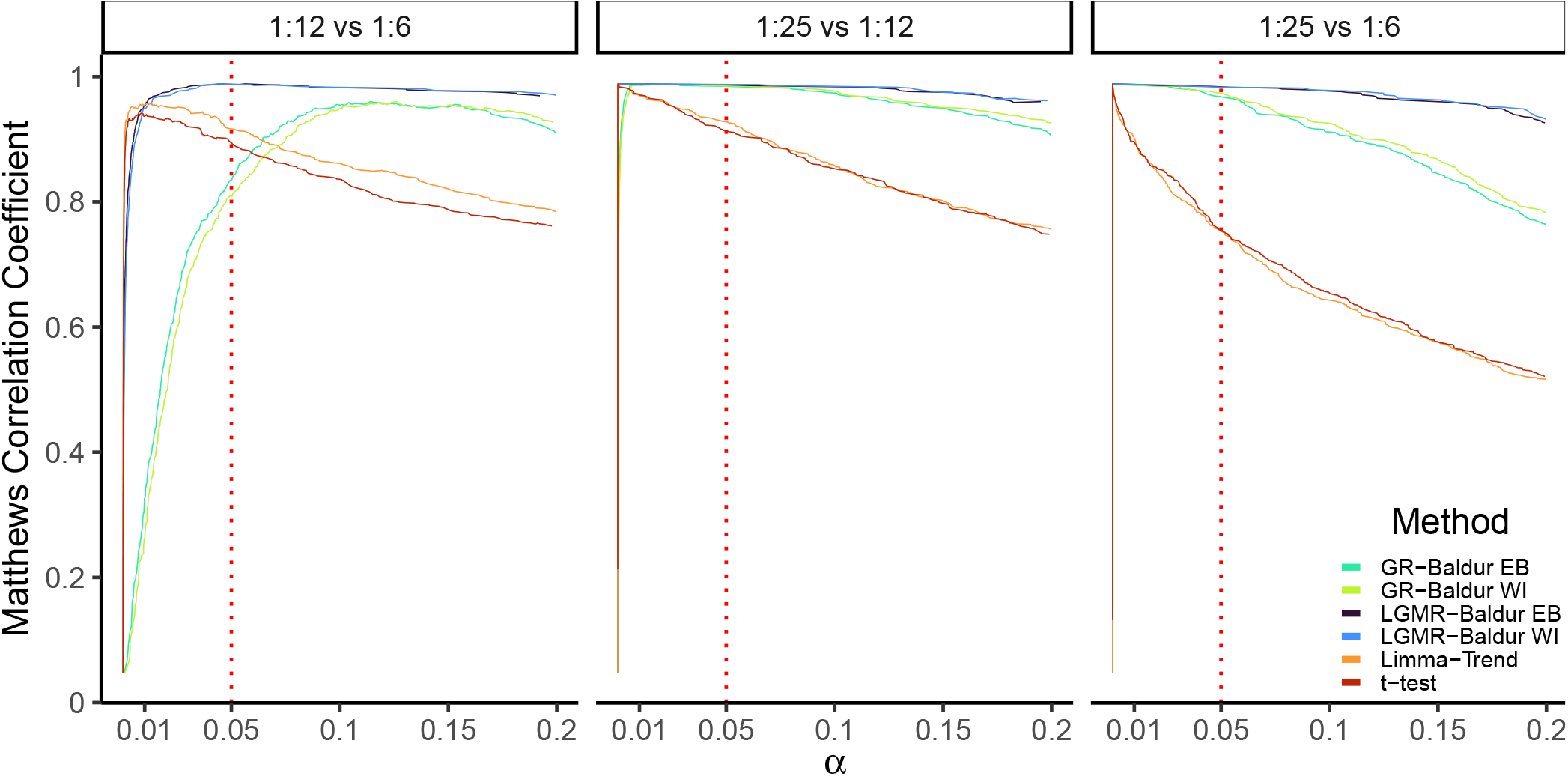
Mathews correlation coefficient of the Human-DS plotted against the significance level (*α*). As in Figure S5.

**Fig. S7.**
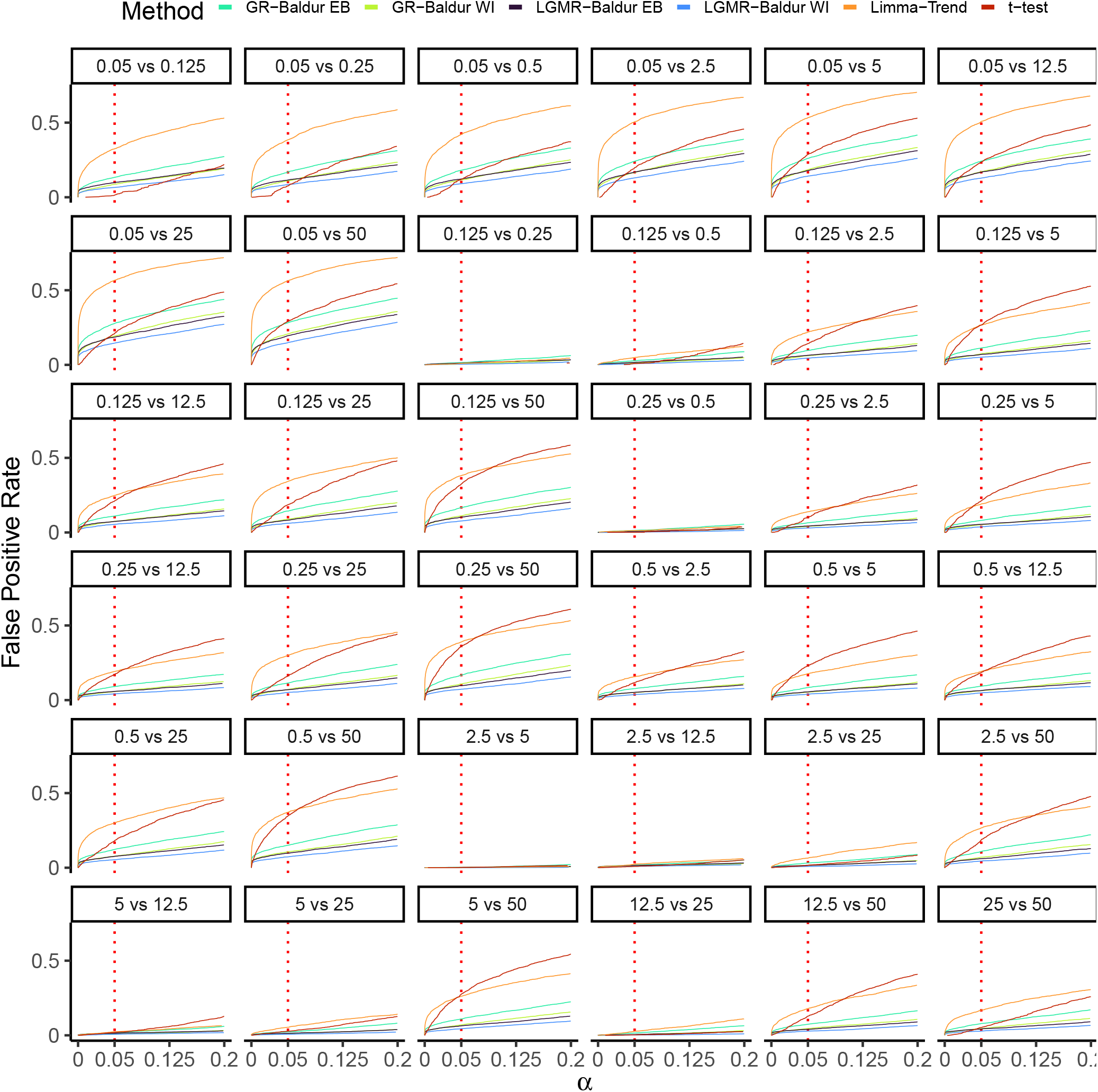
False positive rates of the Ramus-DS plotted against the significance level (*α*). Y-axis shows the false positive rate, X-axis shows the significance level for the different comparisons (as indicated by the box titles), and colors as according to Figure 2.

**Fig. S8.**
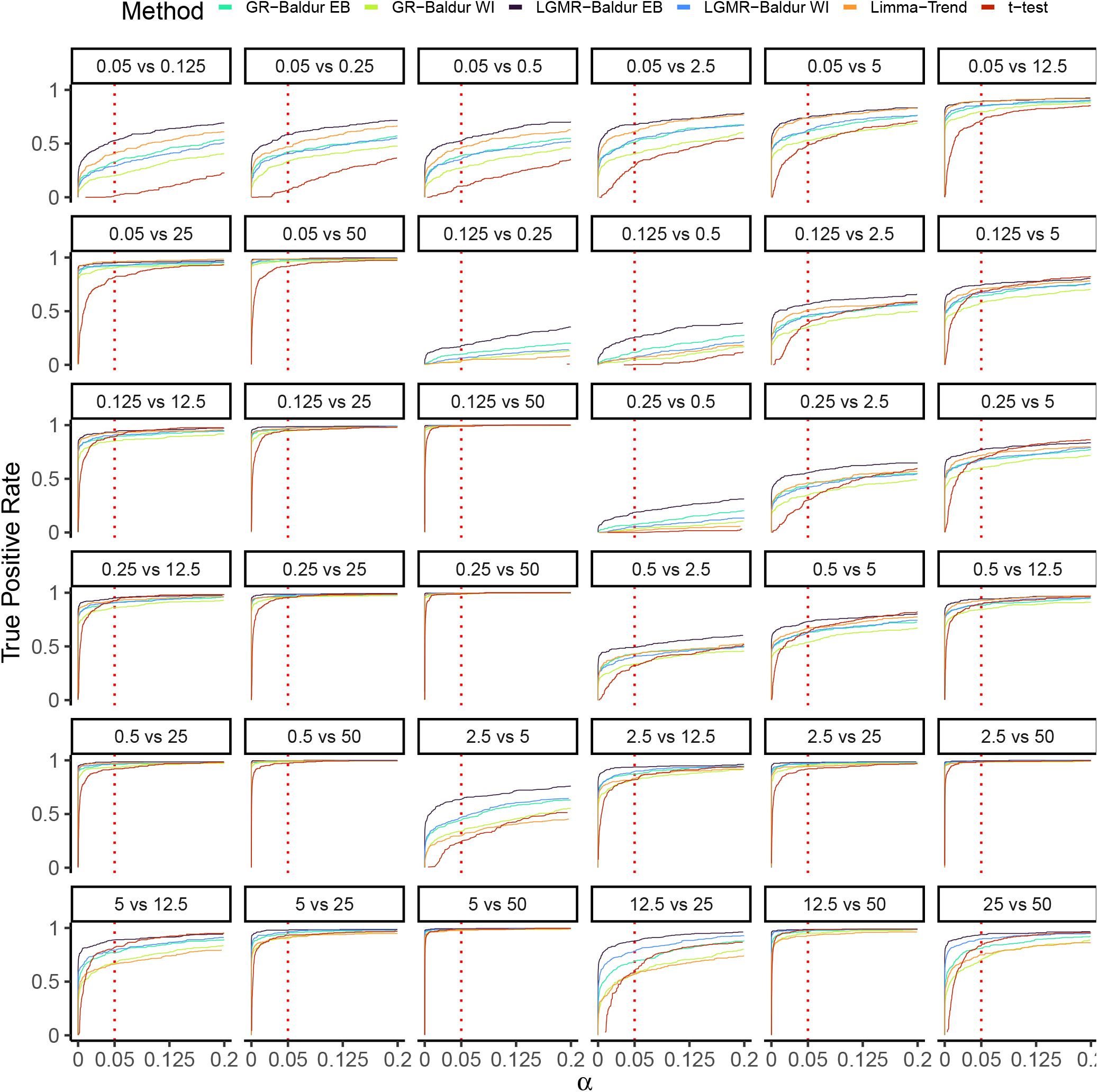
True positive rates of the Ramus-DS plotted against the significance level (*α*). Y-axis shows the true positive rate else as Figure S7.

**Fig. S9.**
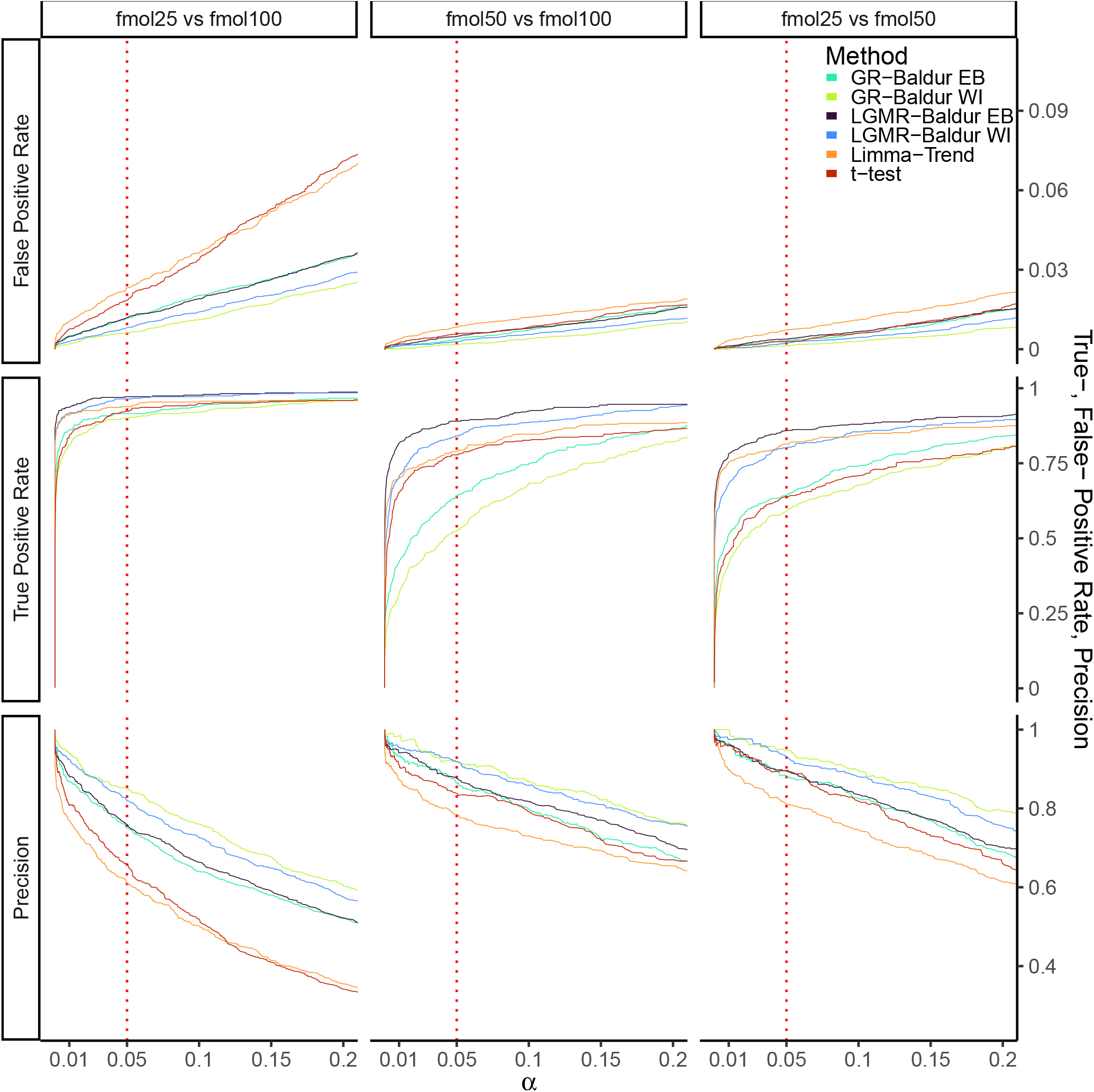
Performance metrics of the pairwise comparisons in the UPS-DS plotted against the significance level (*α*). Y-axis shows the metric value (as indicated by the X-axis facet titles), X-axis shows the significance level for the different comparisons (as indicated by the Y-axis facet titles), and colors as according to Figure 2.

**Fig. S10.**
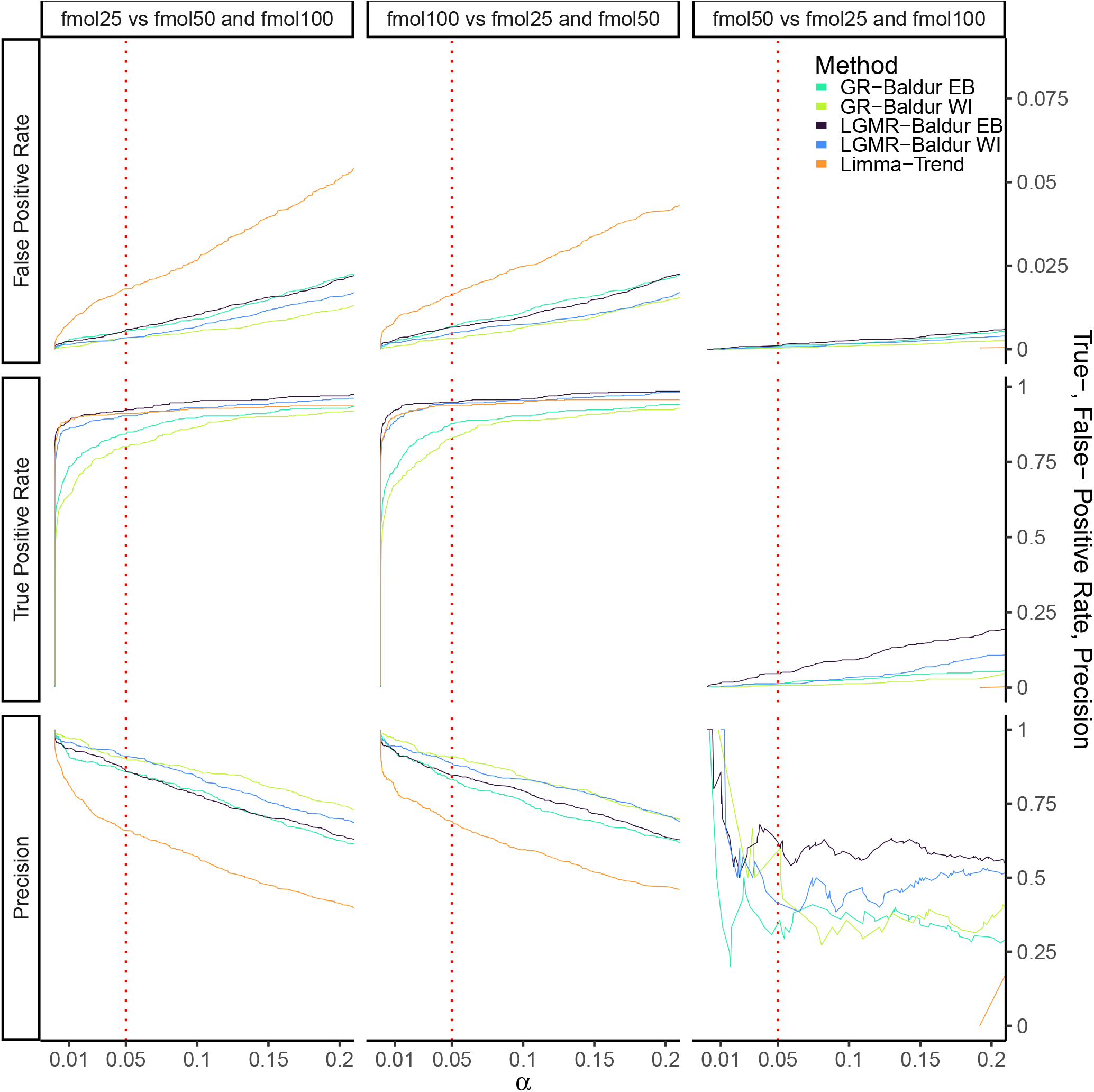
Performance metrics of the complex contrasts in the UPS-DS plotted against the significance level (*α*). Y-axis shows the metric value (as indicated by the X-axis facet titles), X-axis shows the significance level for the different comparisons (as indicated by the Y-axis facet titles), and colors as according to Figure 2. “and” indicates the mean of the two conditions (e.g., contrast vector [-1 0.5 0.5]^T^).

**Fig. S11.**
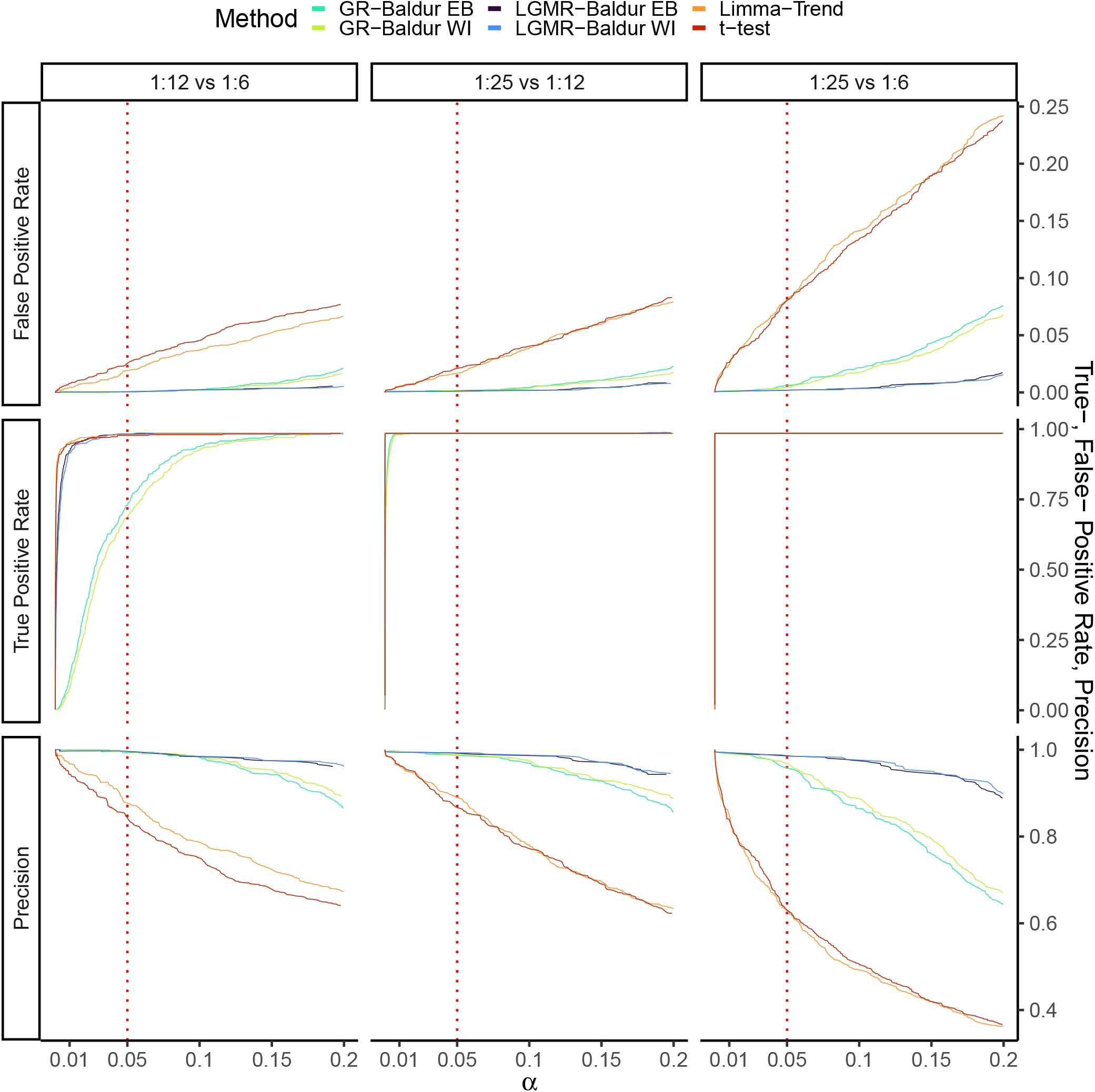
Performance metrics of the Human-DS plotted against the significance level (*α*). As in Figure S9.

**Fig. S12.**
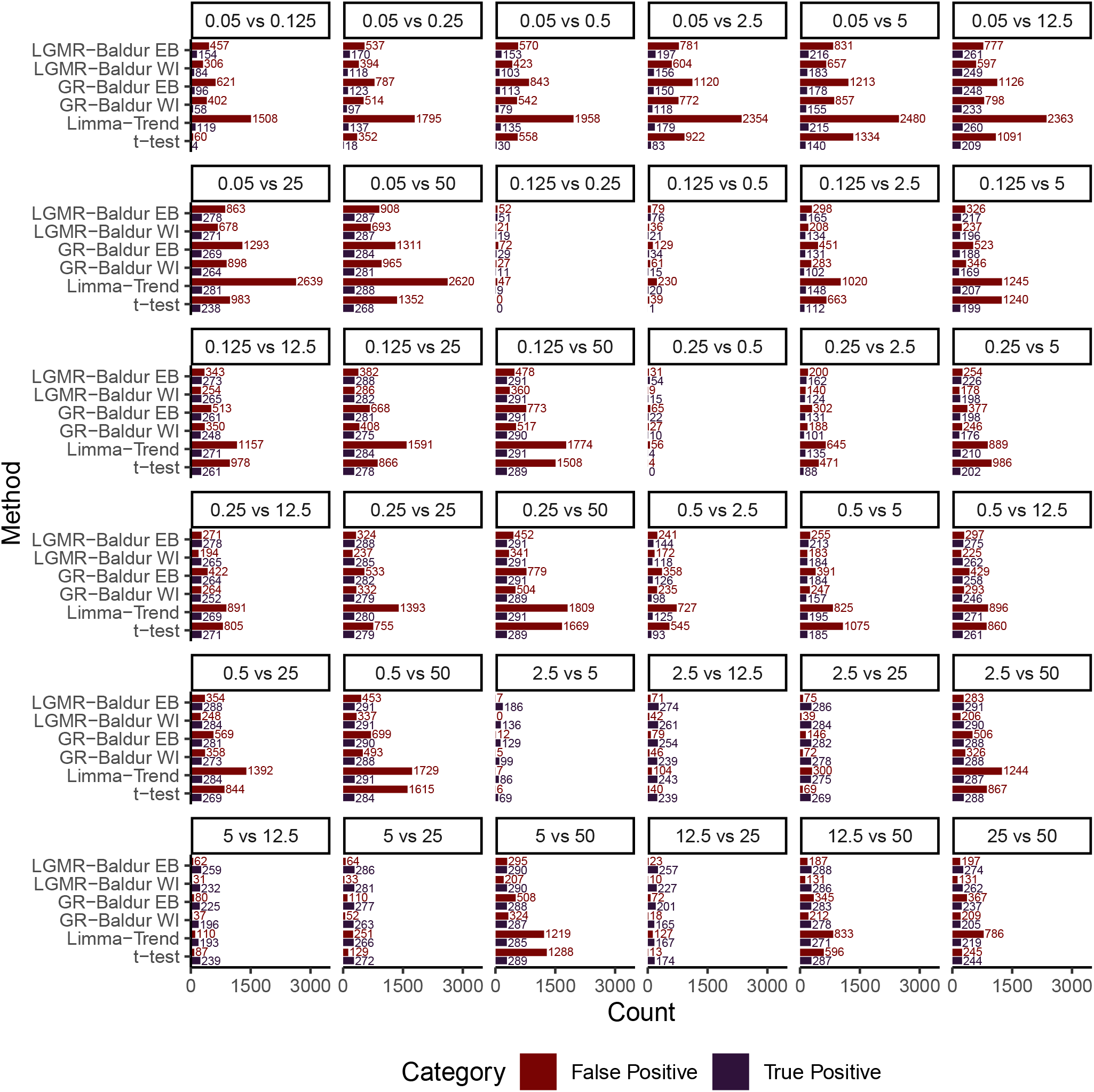
Decision of False Positives and True Positives of at 5 % significance level for the Ramus-DS and methods evaluated here.

**Fig. S13.**
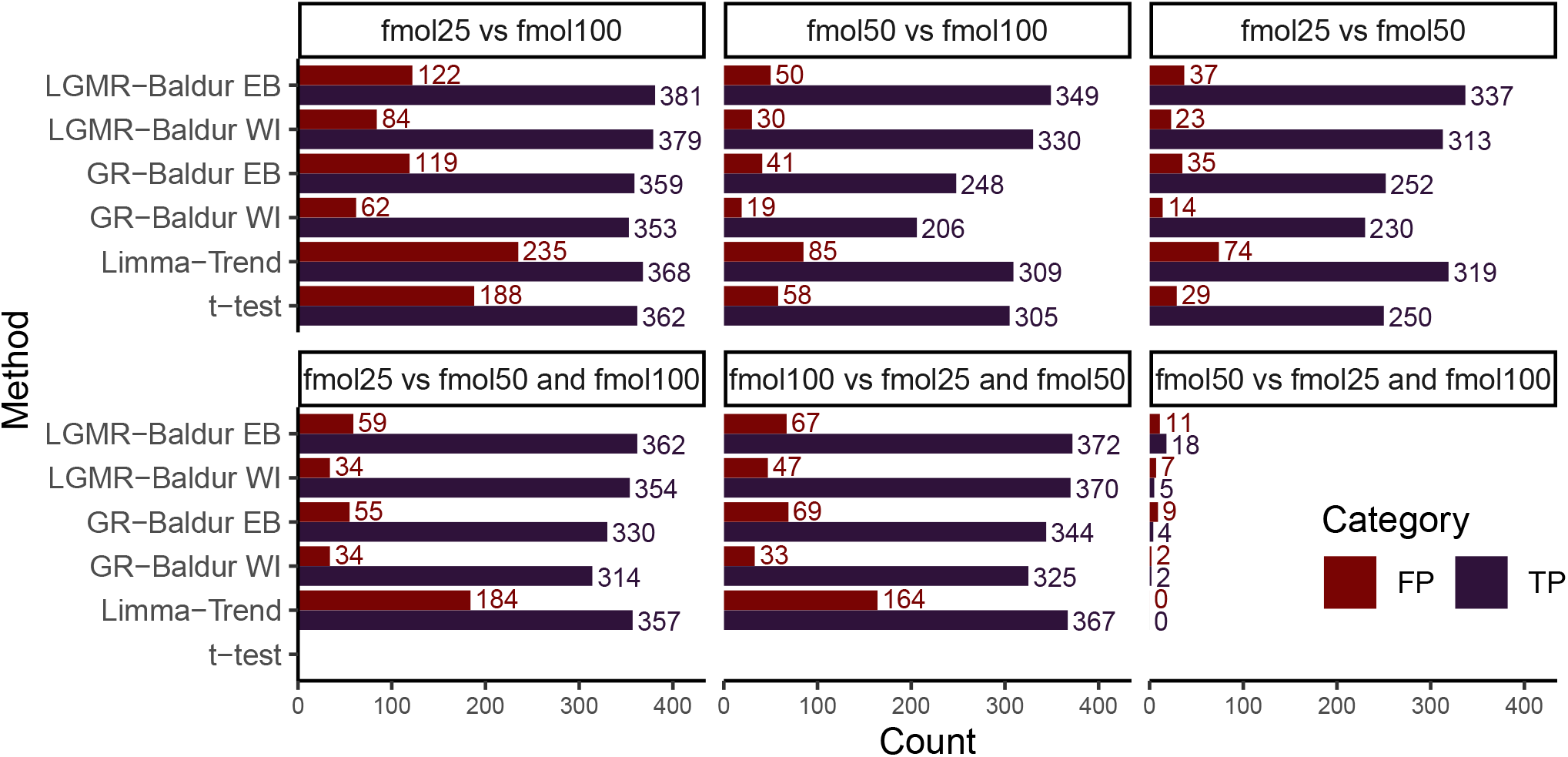
Decision of False Positives and True Positives of at 5 % significance level for the UPS-DS and methods evaluated here. “and” indicates the mean of the two conditions.

**Fig. S14.**
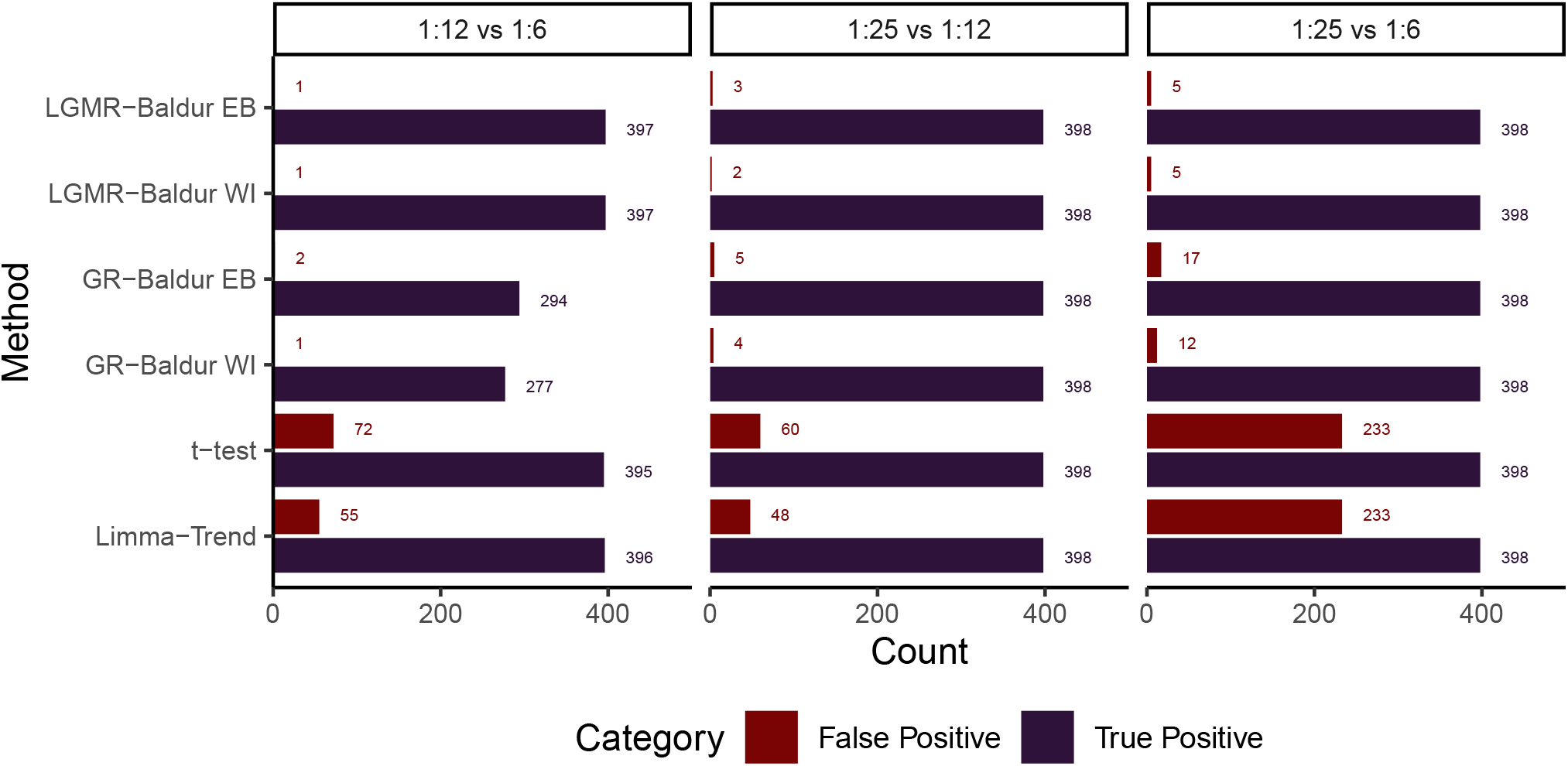
Decision of False Positives and True Positives of at 5 % significance level for the Human-DS and methods evaluated here.

